# Single-cell analysis reveals diverse stromal subsets associated with immune evasion in triple-negative breast cancer

**DOI:** 10.1101/2020.06.04.135327

**Authors:** Sunny Z. Wu, Daniel L. Roden, Chenfei Wang, Holly Holliday, Kate Harvey, Aurélie S. Cazet, Kendelle J. Murphy, Brooke Pereira, Ghamdan Al-Eryani, Nenad Bartonicek, Rui Hou, James R. Torpy, Simon Junankar, Chia-Ling Chan, Eric Lam, Mun N. Hui, Laurence Gluch, Jane Beith, Andrew Parker, Elizabeth Robbins, Davendra Segara, Cindy Mak, Caroline Cooper, Sanjay Warrier, Alistair Forrest, Joseph Powell, Sandra O’Toole, Thomas R. Cox, Paul Timpson, Elgene Lim, X. Shirley Liu, Alexander Swarbrick

**Affiliations:** The Kinghorn Cancer Centre and Cancer Research Division, Garvan Institute of Medical Research, Darlinghurst, NSW 2010, Australia; St Vincent’s Clinical School, Faculty of Medicine, UNSW Sydney, NSW 2052, Australia; Department of Data Sciences, Center for Functional Cancer Epigenetics, Dana-Farber Cancer Institute, Harvard T.H. Chan School of Public Health; Harry Perkins Institute of Medical Research, QEII Medical Centre and Centre for Medical Research, The University of Western Australia, Nedlands, Perth, WA 6009, Australia; Chris O’Brien Lifehouse, Camperdown, NSW 2050, Australia; The Strathfield Breast Centre, Strathfield, NSW 2135, Australia; St Vincent’s Hospital, Darlinghurst, NSW 2010, Australia; Royal Prince Alfred Hospital, Camperdown, NSW 2050, Australia; Pathology Queensland, Princess Alexandra Hospital, Brisbane, Queensland 4102, Australia; Southside Clinical Unit, Faculty of Medicine, University of Queensland, Brisbane, Queensland 4102, Australia; Department of Breast Surgery, Chris O’Brien Lifehouse, NSW 2050, Australia; Royal Prince Alfred Institute of Academic Surgery, Sydney University; RIKEN Center for Integrative Medical Sciences, Yokohama, 230-0045 Japan; Garvan-Weizmann Centre for Cellular Genomics, Garvan Institute of Medical Research, Sydney, Australia; UNSW Cellular Genomics Futures Institute, University of New South Wales, Sydney, Australia; Australian Clinical Laboratories, Northern Beaches Hospital, Frenchs Forest, NSW 2086, Australia

**Keywords:** Cancer associated fibroblasts, single cell RNA sequencing, stromal heterogeneity, triple negative breast cancer, tumour microenvironment

## Abstract

The tumour stroma regulates nearly all stages of carcinogenesis. Stromal heterogeneity in human triple-negative breast cancers (TNBCs) remains poorly understood, limiting the development of stromal-targeted therapies. Single cell RNA-sequencing of five TNBCs revealed two cancer-associated fibroblast (CAF) and two perivascular-like (PVL) subpopulations. CAFs clustered into two states, the first with features of myofibroblasts and the second characterised by high expression of growth factors and immunomodulatory molecules. PVL cells clustered into two states consistent with a differentiated and immature phenotype. We showed that these stromal states have distinct morphologies, spatial relationships and functional properties in regulating the extracellular matrix. Using cell-signalling predictions, we provide evidence that stromal-immune crosstalk acts *via* a diverse array of immunoregulatory molecules. Importantly, the investigation of gene signatures from inflammatory-CAFs and differentiated-PVL cells in independent TNBC patient cohorts revealed strong associations with cytotoxic T-cell dysfunction and exclusion, respectively. Such insights present promising candidates to further investigate for new therapeutic strategies in the treatment of TNBCs.

## Introduction

Heterotypic interactions between stromal, immune and malignant epithelial cells play important roles in solid tumour progression and therapeutic response. Cancer-associated fibroblasts (CAFs) play an integral part in the tumour microenvironment (TME), and can influence many aspects of carcinogenesis including extracellular matrix (ECM) remodelling, angiogenesis, cancer cell proliferation, invasion, inflammation, metabolic reprogramming and metastasis [1]. Recent studies have described roles for CAFs in mediating immune suppression and chemo-resistance, establishing CAFs as novel and attractive targets for anti-cancer therapies in advanced breast cancer [2–6]. Despite their well-described roles in cancer biology, CAFs remain enigmatic: limited studies suggest phenotypic heterogeneity, plasticity and functional diversity, with both tumour-promoting and tumour-suppressive properties [1]. The multi-faceted nature of CAFs suggests that they are comprised of diverse subpopulations, and an improved understanding of stromal heterogeneity may explain how CAFs contribute to the dynamic complexity and functional malleability of the tumour ecosystem.

CAFs of the tumour parenchyma are routinely studied using a handful of markers including α-smooth muscle actin (α-SMA), fibroblast activation protein (FAP), CD90 (THY-1), platelet derived growth factor receptor α and β (PDGFRα and PDGFRβ), podoplanin (PDPN) and fibroblast specific protein 1 (FSP-1, also named S100A4) [1, 7–9]. However, these markers are not necessarily co-expressed, nor specific to the fibroblast lineage [4]. For instance, α-SMA not only identifies CAFs with a myofibroblast morphology but also serves as a general marker for myoepithelial cells and perivascular cells. α-SMA+ cells in the breast tumour stroma can also arise from different mesenchymal lineages including resident fibroblasts, smooth muscle cells and pericytes [10]. In addition, FSP1 is also expressed in macrophages, other immune cells and even cancer cells [11]. Thus, a categorical definition of cancer associated stromal cells and specific cell surface markers remains challenging and is urgently needed [1].

Three broad CAF subtypes have been recently profiled in mouse models of pancreatic ductal adenocarcinoma (PDAC) [12–14]. These are characterised by a myofibroblast-like (myCAFs) phenotype, inflammatory properties (iCAFs) and antigen presenting capabilities (apCAFs) [12–14]. Although little is known about the mechanistic role and clinical relevance of iCAFs and apCAFs, an accumulation of the myCAF marker α-SMA has been shown to correlate with poor outcome in breast and pancreatic cancer [15, 16]. We have shown that targeting Hedgehog-activated CAFs, which have a myofibroblast-like phenotype in ECM regulation, results in markedly improved survival, chemosensitivity and reduced metastatic burden in pre-clinical models of TNBC [3]. In addition, myofibroblast-like CAFs have been shown to contribute to an immunosuppressive microenvironment by attracting T-regulatory cells in breast and ovarian cancer [4, 5]. While these studies point towards the therapeutic targeting of myofibroblast-like CAFs, genetic ablation of α-SMA+ cells in a mouse model of PDAC resulted in more aggressive tumours and reduced mouse overall survival, indicating complex stromal functionalities across distinct tissue sites [17].

Recent advances in single-cell RNA sequencing (scRNA-Seq) have overcome some of the technical hurdles in the investigation of cellular heterogeneity amongst complex tissues such as carcinomas. Recent patient studies have dissected the TME in head and neck squamous cell carcinomas and lung tumours, revealing new insights into stromal and immune subsets associated with disease progression [18, 19]. Single-cell studies of human breast cancers have been limited to immune cells, while studies in mouse models have revealed four subclasses of CAFs [20]. Although CAFs from human breast carcinomas have been profiled by flow cytometry and bulk sequencing, comprehensive single-cell profiling has yet to be performed in TNBC patients [4].

TNBC is an aggressive breast cancer subtype, which is lacking in effective targeted therapeutic options. It is clinically defined by negative status for targetable hormone receptors (estrogen receptor and progesterone receptor) or HER2 amplification. Studies in mice and humans have demonstrated that TNBC progression can be influenced by stromal cells however, a comprehensive understanding of the stromal hierarchy is yet to be established [2–6]. To investigate this in more detail, we performed unbiased high-throughput scRNA-Seq to profile the TME directly in patient tumour tissues. In addition to CAFs, we identified stromal cells with a perivascular-like (PVL) profile, which were not necessarily associated with blood vessels. Our study focuses exclusively on CAFs and PVL cells, which we collectively refer to as ‘stroma’. Using orthogonal methods, we found that functions previously ascribed to CAFs as unitary cell types are actually performed by specialised subsets of stromal cells with distinct morphological, spatial and functional properties [20]. In addition, by sampling cells from the entire TME, we were able to predict paracrine signalling between stromal and immune cell subsets. From this, we analysed large patient gene expression datasets to show significant association between inflammatory-like CAFs and differentiated-PVL cells with immune evasion. Our human TNBC single-cell datasets provide a new taxonomy of human cancer-associated stromal cells, which we envisage can be used to further develop TME-directed therapies.

## Results

### Composition of triple-negative breast cancers at cellular resolution

We performed scRNA-Seq on primary breast tumours collected from five patients (Fig. EV1A-B) using a marker free approach. Fresh tissues were dissociated into single cell suspensions prior to single-cell capture on the Chromium controller (10X Genomics) and sequencing on the NextSeq 500 (Illumina) (Fig. 1A; Fig. EV1C). In total, we sequenced 24,271 cells, with an average of 4,854 cells per patient (Fig. EV1D). A total of 28,118 genes were detected with an average of 1,658 genes expressed, and 6,215 unique molecular identifiers (UMIs) detected per cell (Fig. EV1E-H). Data from individual tumours were integrated and clustered using canonical correlation analysis (CCA) in Seurat [21].

**Figure 1.**
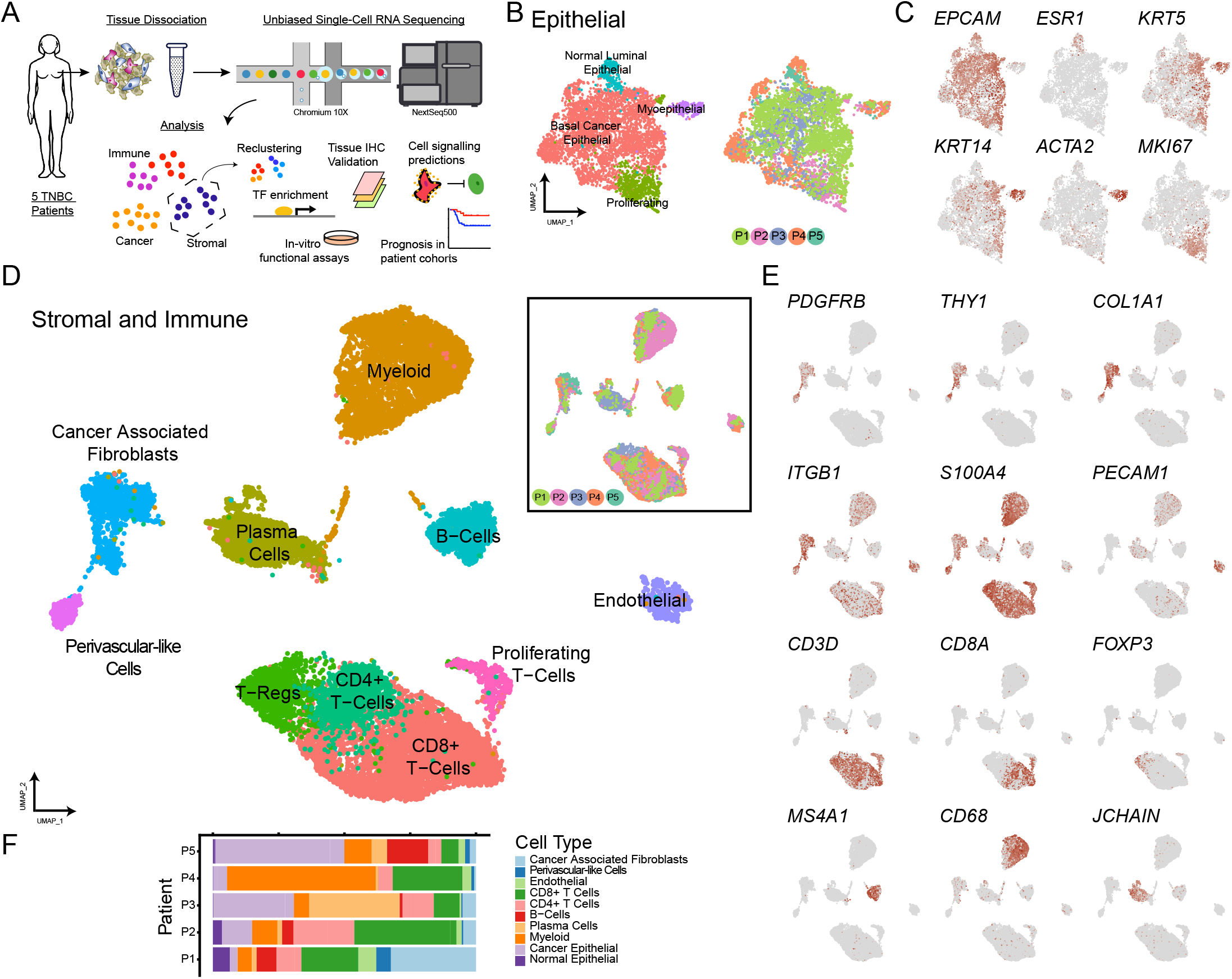
Cellular composition of five triple-negative breast carcinomas. **a,** Schematic highlighting the application of our single-cell RNA sequencing experimental and analytical workflow for primary patient tissue. **b,** UMAP visualisation of 4,986 epithelial cells aligned using canonical correlation analysis in Seurat. Cells are coloured by their cell type annotation (left) and patient of origin (right). **c,** Log normalised expression of markers for epithelial (*EPCAM*), mature luminal epithelial (*ESR1*), myoepithelial (*KRT5, KRT14* and *ACTA2*) and proliferating cancer cells (*MKI67*). **d,** UMAP visualisation of 19,285 stromal and immune cells aligned and visualised as represented in **b**. ***e***, Log normalised expression of markers for fibroblasts (*PDGFRB*, *THY1*, *COL1A1*, *ITGB1* and *S100A4*), endothelial cells (*PECAM1*), T-cells (*CD3D*), CD8 T cells (*CD8A*), T-regulatory cells (*FOXP3*), B-cells (*MS4A1*), myeloid cells (*CD68*) and plasma cells (*JCHAIN*). **f,** Proportion of cell types across each patient.

Epithelial cells (Fig. 1B-C) and stromal-immune cells (Fig 1D-E) were first annotated through the expression of canonical cell type gene markers. This revealed four major cell states within the epithelial compartment (Fig. 1B-C), including a major cluster of 4,095 cancer cells (16.9% of all cells; *EPCAM*^+^, *ESR1*^*−*^) and a second cluster of 614 cancer cells with high proliferation (2.5%; *MKI67*^*+*^). The remaining two smaller epithelial clusters had gene expression features consistent with normal luminal (277 cells, 0.9%; *EPCAM*^+^, *ESR1^+^*) and myoepithelial cells (212 cells, 0.9%; *EPCAM*^lo^, *KRT5^+^, KRT14^+^* and *ACTA2^+^*). Neoplastic or normal status of these cell clusters was confirmed by inferring genome copy number alterations over large genomic regions using InferCNV (Appendix Fig. S1) [22]. In addition to marker genes, stromal and immune clusters were further classified through scoring against published cell type signatures from the XCell database with an area under the curve approach (AUCell) (Fig. EV2A) [23, 24]. In the immune compartment (Fig 1D-E), we identified 7,990 T-lymphocytes (32.9%; *CD3D*), 1,245 B-cells (5.1%; *MS4A1*), 1,955 plasma cells (8.1%; *JCHAIN*) and 4,606 myeloid cells (19.0%; *CD68*). Through re-clustering of the T-lymphocytes (Fig. EV2B-D), we identified 175 T-follicular helper cells (2.2%; *CXCL13* and *CD200*), 994 T-Regulatory cells (12.4%; *FOXP3* and *BATF*), 2,003 other CD4+ T-cells (25.1% of all T-cells; *CD4, IL7R* and *CD40LG*), 3,691 CD8+ T-cells (46.2%; *CD8A* and *GZMH*), 605 proliferating T-cells (7.6%; *MKI67*), 358 NK Cells (4.5%; *GNLY*, *KLRD1*, *NCR1, XCL1 and NCAM1*) and 164 NKT cells (2.1%; *GNLY, KLRD1, NCR1 and CD3D*^*−*^). The remaining cells consisted of 610 endothelial cells (2.5%; *PECAM1*) and two distinct clusters (with 1,409 and 320 cells, 5.8% and 1.3%, respectively) sharing the expression of common stromal markers including *PDGFRB*, *S100A4* (FSP-1), *ITGB1* (CD29) and *THY1* (CD90). These non-endothelial nor immune cells (collectively referred to as stromal in this study) were enriched for a fibroblast cell type signature from XCell (Fibroblasts_FANTOM_1; Fig. EV2A). All annotated cell types were detected in each patient, with varying proportions of cell types between cases, indicating no patient specific sub-populations in our integrated dataset (Fig. 1F).

### Reclustering stromal cells revealed four distinct sub-clusters in human TNBCs

Although the stromal clusters shared many common markers used to study CAFs, we further inspected their heterogeneity through reclustering each population (Fig. 2A). Sub-clusters were detected across multiple clustering resolutions in the *FindClusters* function in Seurat (resolutions 0.2, 0.3 and 0.4), with varying proportions from each patient (Fig. 2B). The first cluster, which was classified as CAFs through the expression of fibroblast-specific markers *(PDGFRA, COL1A1, FAP* and *PDPN*), formed two sub-clusters (Fig. 2A-C). The first CAF sub-cluster was comprised of 280 cells (16.2% of all stromal; red cluster) and was classified as myofibroblast-like CAFs (myCAFs) through the elevated expression of activated fibroblast markers (*ACTA2, FAP* and *PDPN)* and collagen-related genes (*COL1A1* and *COL1A2*) (Fig. 2C-D) [12–14]. The second CAF sub-cluster comprised of 1,129 cells (65.3%; orange cluster; Fig. 2A-C) and resembled inflammatory-CAFs (iCAFs) through the enrichment of the CAF chemokine marker *CXCL12* (also known as SDF-1) (Fig. 2C-D) [12–14]. We next compared our CAF clusters to the subsets previously reported in pancreatic cancer [12–14]. This was performed by scoring published CAF gene signatures across our stromal clusters using the AUCell method (Fig. EV2E) [23]. This revealed the enrichment of pancreatic myCAF and iCAF signatures in our breast myCAF and iCAF clusters, respectively, suggesting similar phenotypes likely exist across both tissue sites (Fig. EV2E). While the signatures were largely conserved, a number of human PDAC CAF markers were detected in opposing cell types, for example *IL6* was expressed by PVL cells rather than iCAFs (Fig. EV2F). No clusters showed any particular enrichment for signatures of antigen-presenting CAFs, potentially because they are a rare cell type that was not sampled, or are unique to pancreas tumours (Fig. EV2E).

**Figure 2.**
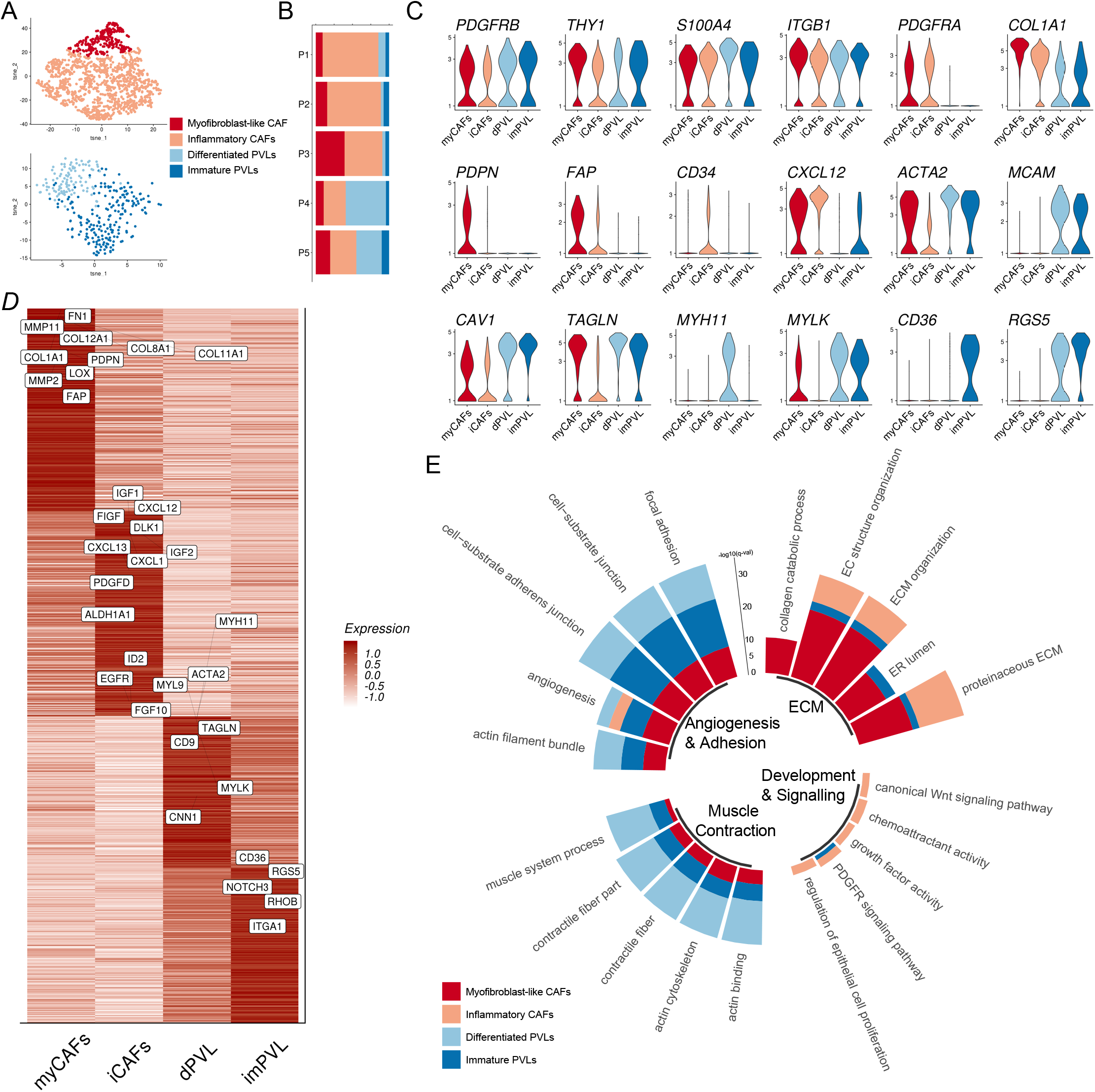
Stromal landscape of TNBCs reveals four subpopulations of cancer-associated fibroblasts and perivascular-like cells. **a,** *t-*SNE representation of the four subclasses of cancer-associated fibroblasts (CAFs) and perivascular-like cells (PVL), named myofibroblast-like CAFs (myCAFs; 280 cells), inflammatory-like CAFs (iCAFs; 1,129 cells), differentiated-PVL cells (dPVL cells; 122 cells) and immature-PVL cells (imPVL cells; 198 cells). **b,** Plot showing the composition of the four stromal subsets across all five patients. **c,** Expression of parenchymal markers commonly associated with CAFs and perivascular cells. **d,** Cluster averaged log normalised expression of the top 300 differentially expressed genes between the four stromal subsets with stromal-related genes of interest annotated. Expression values are scaled per cluster. **e,** Circle histogram plot of the top gene-ontologies enriched in each of the four stromal subsets, with pathways broadly grouped for ECM, development and signalling, muscle contractile-features and angiogenesis and adhesion. Scale bar represents the −log10 q-value for the enrichment of individual GO terms, as determined using ClusterProfiler.

In contrast, the second stromal cluster was enriched for perivascular markers, including genes associated with pericytes and smooth muscle cells (*ACTA2, MCAM, CAV1, TAGLN, MYH11, MYLK* and *RGS5;* Fig. 2C-D) [25]. *MCAM* (also known as CD146) has shown to be a robust marker to differentiate perivascular cells from fibroblasts in human tissues [26–29]. PVL cells were further classified as either differentiated-PVL (dPVL; 122 cells in light blue, 7.1%), characterised through the enrichment of myogenic differentiation genes (*TAGLN*, *MYH11* and *MYLK*), or immature-PVL (imPVL; 198 cells in dark blue; 11.5%), characterised by the elevated expression of genes associated with an immature phenotype (*PDGFRB, CD36* and *RGS5)* (Fig. 2C-D) [30]. To our surprise, both PVL subsets were also enriched for the human PDAC myCAF signature, suggesting PVL cells share some similarities in gene expression profile with myCAFs (Fig. EV2E-F). All four stromal subsets were detected in all five patients, however there were differences in the proportions between the patients (Fig. 2B; Fig. EV3A-B). The stromal profiles of Patient-1 (P1) and P2 were predominantly comprised of iCAFs, myCAFs were highest in P3, whilst PVL cells were highly abundant in P4 and P5 (Fig. 2B; Fig. EV3A-B).

Next, we identified differentially expressed genes (DEGs) between the four subsets using the MAST method, which compares each subset against all other subsets [31]. This identified a total of 894, 610, 258 and 289 DEGs (log fold change threshold of 0.1, *p-value* threshold of 1×10^−5^ and FDR threshold of 0.05) by myCAFs, iCAFs, dPVL and imPVL cells, respectively (Fig. 2D; Appendix Table S1). We performed gene ontology (GO) analysis using the top 250 DEGs from each subset using the clusterProfiler tool (Fig. 2E; Appendix Table S2) to determine the pathway level differences driving stromal heterogeneity [32]. This revealed an enrichment of collagen biosynthesis and ECM-regulatory pathways in myCAFs, which included fibrillar collagen genes *COL1A1* and *COL1A2* and ECM remodelling metalloproteinases *MMP1* and *MMP11* (Fig. 2D-E). We identified the enrichment of developmental signalling pathways and chemotactic regulation in iCAFs, including soluble factors such as *IGF1, FIGF* and *PDGFD,* and the chemokines *CXCL12* and *CXCL13* (Fig. 2D-E). Stem cell markers including *ALDH1A1* and *ID2,* and the growth factor receptor *EGFR* were also upregulated in iCAFs (Fig. 2D). Within the PVL cells, the dPVL cluster was enriched for pathways related to the muscle system and contractility, while the imPVL cluster was enriched for pathways related to focal and substrate adhesion, including the integrin molecule *ITGA1* (Fig. 2D-E). No stromal clusters expressed canonical markers for proliferation, including *MKI67* and *AURKA*. As many of the genes and pathways identified were related to cell activation and contractility, we hypothesised that the stromal sub-clusters resembled cell differentiation stages rather than distinct subpopulations. Cell trajectories were examined using the Monocle method, which revealed subsets of CAFs and PVL cells distributed across pseudotemporal space (Fig. EV3C-D) [33]. For example, *COL1A1, ACTA2* and *CXCL12* expression transitioned throughout CAF differentiation (Fig. EV3C), while *CD36*, *RGS5* and *MYH11* transitioned throughout PVL differentiation (Fig. EV3D). Our findings indicate that the stroma in TNBC is comprised of four major transcriptional states related to cell differentiation, which branch from the two major fibroblast and perivascular-like lineages.

### Transcription factor pathways enriched across stromal subclasses

We next sought to investigate if gene regulatory networks could further explain the underlying heterogeneity in stromal subpopulations. To examine the activity of CAF and PVL transcription factors (TFs), we applied the SCENIC method to build gene regulatory networks from scRNA-Seq data and identify activating cis-regulatory elements [23, 34]. Through applying this to the normalised stromal gene expression matrix, SCENIC identified a total of 190 activated TFs, of which 166 were identified to be significantly different across the four stromal subsets (one-way ANOVA; *p-value* threshold of 1×10^−5^). We focused on the top 50 strongest candidates based on their average AUC values (Fig. 3; Appendix Fig. S2).

**Figure 3.**
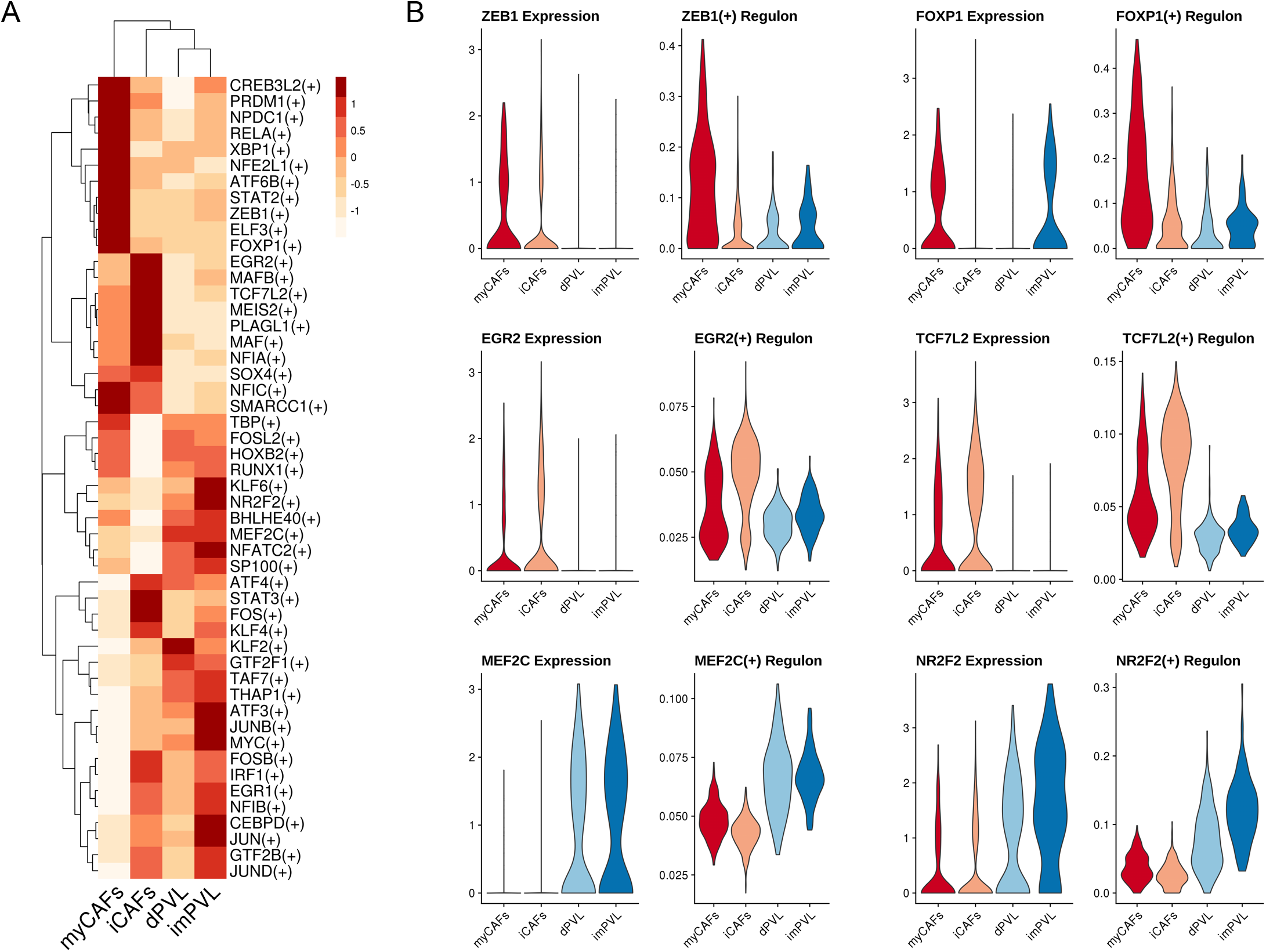
Polarised gene regulatory states between cancer-associated fibroblasts and perivascular-like subclasses. **a,** Polarised gene regulatory states underlying stromal subclasses. Heatmap shows the averaged regulon activity (area under the curve; AUC) for the top 50 highest TFs regulons as estimated using SCENIC. All regulons are statistically enriched across the four subsets (p < 1×10^−5^ One-way ANOVA). Heatmap is clustered using Euclidean distance and complete linkage. **b,** Candidate transcriptional drivers of each CAF and PVL subset. Violin plots showing the log normalised gene expression (left) of the TF and its respective AUC regulon activity (right). TFs ZEB1 and FOXP1 enriched in myofibroblast-like CAFs, EGR2 and TCF7L2 enriched in inflammatory-like CAFs, MEF2C enriched in PVL cells and NR2F2 enriched in immature-PVL cells.

In examining the top candidate TFs (Fig. 3; Appendix Fig. S2), *ZEB1* and *FOXP1* were enriched in myCAFs. A recent study inhibiting stromal ZEB1 in the PyMT mouse model of breast cancer reduced tumour growth, invasion and impaired ECM deposition [35]. In other tissue contexts, FOXP1 was reported to regulate the fibrotic potential of stromal cells via the Wnt/beta-catenin pathway, including myCAF marker genes such as *ACTA2* and *COL1A1* [36]. Known roles of such TFs are consistent with the predicted ECM-regulating phenotype of myCAFs. The *EGR2* and *TCF7L2* regulons were enriched in iCAFs (Fig. 3). EGR2 is known to regulate the expression of immunomodulatory molecules in mesenchymal stem cells [37]. The TCF family including TCF7L2 (also known as TCF4) are Wnt-regulated TFs that are highly expressed during early development [38]. As iCAFs also expressed the stem cell markers *ALDH1A1* and *ID2*, we hypothesised that they resemble a stem or progenitor-like state.

For PVL cells, *MEF2C* was a highly enriched driver in both subsets (Fig. 3). Myocyte enhancer factor 2 (MEF2) is a well-defined regulator for the development of vascular smooth muscle cells [39, 40]. We identified *KLF2* enriched in dPVL cells, and *NR2F2* enriched in imPVL cells (Fig. 3). KLF2 is required for smooth muscle cell migration and maturation in blood vessel formation, consistent with the predicted differentiation state of dPVL cells [41]. Furthermore, NR2F2, also known as COUP-TFII, is highly expressed by myogenic precursors and is known to inhibit muscle development, which is consistent with the predicted immature state of imPVL cells [42]. In summary, we identified unique and novel TF drivers in each of the four stromal subclasses, providing further insights into the transcriptional drivers underlying stromal heterogeneity.

### Validation of stromal subsets in primary breast cancer tissue

To validate the existence of the four stromal subclasses described above in TNBC patient tissue (Fig. 4A), we first performed fluorescence-activated cell sorting (FACS) isolation on scRNA-Seq matched human tissue sections (Fig. 4B). Our gating strategy used EPCAM, CD45 and CD31 as negative markers to exclude epithelial, immune and endothelial cells, respectively (Fig. 4B). We additionally used PDGFRβ to positively select all stromal populations and avoid contaminations from cancer stem cells and breast myoepithelial cells which have low EPCAM expression [43, 44]. Based on our initial scRNA-Seq findings, we determined PDGFRα and CD146 (*MCAM*) as good markers to discriminate CAFs and PVL cells, respectively. Following the initial isolation and culturing of CAFs (PDGFRβ^+^/PDGFRα^+^/CD146^−^) and PVL cells (PDGFRβ^+^/PDGFRα^−^/CD146^+^), we next performed simultaneous FACS analysis of additional stromal markers to validate the presence of the four stromal subsets in culture. We show that myCAFs and iCAFs could be distinguished by FAP^HIGH^/CD90^HIGH^ and FAP^LOW^/CD90^LOW^ expression, respectively (Fig. 4B, Fig. EV3E-F), whilst imPVL cells could be discriminated from dPVL cells by CD36^+^ expression (Fig. 4B, Fig. 4B). We validated the gene expression of cultured bulk and sorted CAF fractions using quantitative real time PCR (qPCR) (Fig. EV3G). As controls, *PDGFRA* and *PDGFRB* were expressed in both the FAP-high and FAP-low populations. Consistent with the FACS sorting strategy and scRNA-Seq findings, *FAP* and *ACTA2* were enriched in FAP^HIGH^(myCAF) sorted cells, while *CXCL12* and *EGFR* were enriched in FAP^LOW^ (iCAF) sorted cells (Fig. EV3G). We next performed immunofluorescence (IF) to further validate additional markers and explore potential morphological differences. Here, α-SMA expression was used to identify myCAFs from iCAFs (Fig. 4C; Fig. EV4A), and CD36 to distinguish imPVL from dPVL cells (Fig. 4D; Fig. EV4B). From our observations, myCAFs and dPVL cells had a more elongated morphology in comparison to iCAFs and imPVL cells (Fig. 4C; Fig. EV4A), which is consistent with the predicted differentiation state of each subset. Importantly, we defined a novel gating strategy that allowed us to purify the four stromal subsets for subsequent *in vitro* functional characterisation.

**Figure 4.**
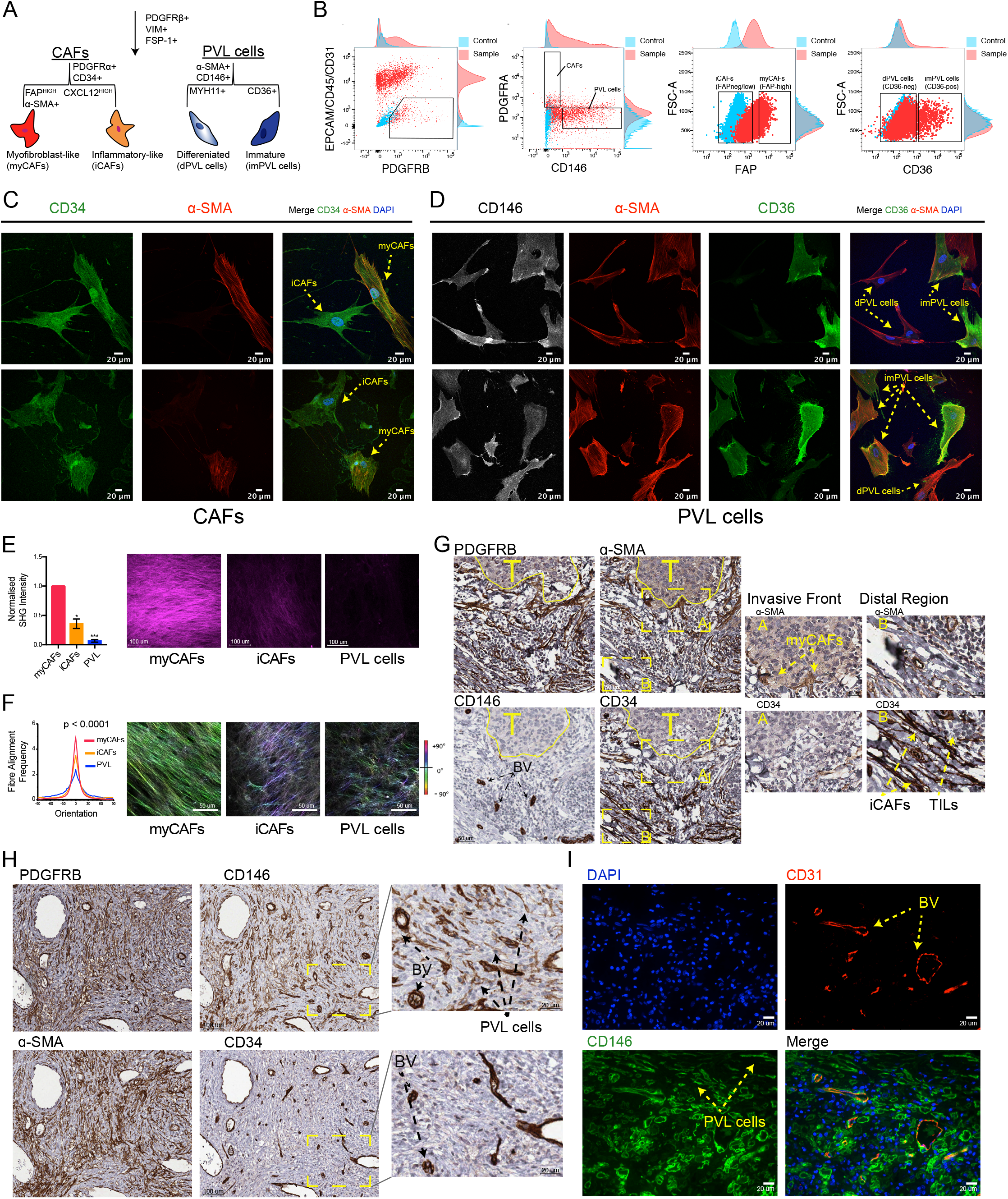
Morphological, phenotypic and spatial differences underlying stromal heterogeneity. **a,** Summary of the markers distinguishing each of the four stromal subpopulations identified in this study. **b,** FACS validation in matched patient tissue. Stromal cells are negatively gated for EPCAM (epithelial), CD45 (immune) and CD31 (endothelium) and positively selected for PDGFRβ. Subsequent markers PDGFRα and CD146 (MCAM) are used to distinguish CAFs and PVL cells, respectively. Expression of FAP^HIGH^, FAP^LOW^, CD36^+^ and CD36^−^ are further used to define myofibroblast-like CAFs, inflammatory-like CAFs, immature-PVL cells and differentiated-PVL cells, respectively. **c-d,** Immunofluorescence of cultured human CAFs **(c)** and PVL cells **(d)**, staining for CD34 (CAFs), α-SMA (myCAFs and PVL cells), CD146 (PVL cells) and CD36 (imPVL cells). **e-f,** Quantitative analysis of collagen abundance **(e)** and orientation **(f)** using second harmonic generation (SHG) from cellular derived matrices from stromal subsets and representative images multiphoton SHG images (n = 3 biological replicates). Statistical significance for collagen abundance **(e)** was determined using unpaired two-tailed Student’s t test with equal standard deviation. After normalization of the orientation peak distributions **(f)**, statistical significant was determined using a Kruskal-Wallis test with Dunn’s post-hoc multiple comparisons test (p value <0.05). **g-h,** Immunohistochemical staining of PDGFRβ, α-SMA, CD34 and CD146 in serial sections cut 4 μm apart from matched cases; Patient-2 **(g)** and Patient-4 **(h).** Images were aligned using FIJI. Co-localisation of CD34 and CD146 was used to distinguish blood vessels, where their differential staining was used to identify CAFs and PVL cells. **g,** MyCAFs were found to be localised at the invasive stromal interface, whilst iCAFs were located at distal regions. **h,** Case with a high abundance of PVL cells in regions surrounded by blood vessels. **i,** Validation of detached PVL cells from blood vessels using co-immunofluorescence of CD31 (red), CD146 (green) and DAPI (blue). Representative images from Patient-4 is shown.

### Myofibroblast-like CAFs have elevated capabilities for collagen secretion and alignment

From the above results, we predicted myCAFs to be the predominant subset synthesising ECM components. To investigate this, we generated cell-derived matrices (CDMs) to compare the ability of each human stromal subset to lay down collagen, as previously described [45]. Purified stromal subsets were seeded and cultured onto glass for 7 days. To assess Collagen I deposition, we used Second Harmonic Generation (SHG) microscopy, which is a sensitive method for quantifying fibrillar collagen density and orientation in an unlabelled manner. This revealed FAP^HIGH^ myCAFs had a significant increase in SHG signal intensity compared to FAP^LOW^ iCAFs, while PVL cells had a significantly lower SHG signal compared to both CAF subsets (Fig. 4E). Higher densities of stromal collagen is a hallmark of breast tumour growth, invasiveness, and risk of disease development [46–48]. Our findings also indicate that PVL cells do not adopt fibroblast-like traits in contributing to the collagenous TME. Further analyses of collagen fibre orientation also revealed that in addition to increased amounts, the orientation of the collagen fibres deposited by myCAFs was more uniformly aligned compared to iCAFs and PVL cells (indicated by the higher, narrow peak in Fig. 4F). It has been previously shown that tumour associated collagen signatures (TACs), characterised by the alignment of collagen fibres, is a good factor for predicting breast cancer survival [49]. In further parallels to pancreatic cancers, FAP-overexpressing fibroblasts have been shown to produce more parallel aligned fibres, enhancing the directionality and velocity of cancer cell invasion [50]. Importantly, these data highlights that the regulation of the ECM, namely in collagen density and orientation, is mainly regulated by the specialised myCAF subsets. In summary, our findings demonstrate that the stromal subclasses described here are functionally distinct, and provide a novel strategy for their purification from breast cancers.

### Stromal subclasses are spatially distinct

To investigate the spatial localisation of CAFs and PVL cells, we performed immunohistochemistry (IHC) with markers identified by scRNA-Seq on data matched patient tissues. We also wanted to validate that CAFs and PVL cells localise to the intratumoural regions of tumour specimens and are not from adjacent normal tissue or blood vessels. We stained serial 4 μm sections and identified stromal cell types using a combination of markers identified previously by scRNA-Seq and DGE (Fig. 2C): pan-stromal (PDGFRβ^+^), myCAFs (PDGFRβ^+^, α-SMA^HIGH^ and CD146^−^), iCAFs (PDGFRβ^+^, α-SMA^−^, CD34^HIGH^ and CD146^−^) and PVL cells (PDGFRβ^+^, α-SMA^HIGH^, CD34^−^ and CD146^+^). As CD34 and CD146 are commonly used markers of the endothelium but are mutually exclusive in CAFs and PVL cells, we used their co-localisation in combination with PDGFRβ staining and morphology (rings surrounding lumen) to identify endothelial cells [26]. This IHC strategy revealed regions where myCAFs (α-SMA^HIGH^) were located in close proximity to the invasive tumour interface, while iCAFs (CD34^HIGH^) were relatively distal to this interface (Fig. 4G). In these particular cases, no PVL cells were present in these regions and CD146 was completely restricted to blood vessels (Fig. 4G). In distal regions which were enriched for iCAFs, we also identified a high co-localisation of tumour-infiltrating lymphocytes as identified by morphology (Fig. 4G).

By definition vascular smooth muscle cells (vSMCs) and pericytes should be localised around arteries and veins to facilitate vascular development and stability. To examine whether PVL cells are vessel-associated, we used co-IF staining for CD31 and CD146 to mark endothelial cells and PVL cells, respectively. We readily detected PVL cells at non-blood vessel regions in the stroma of 4 out of 5 matched patient tissue sections (all cases except P3), including P4 where it was highly abundant (Fig. 4H-I; Fig. EV4C). Consistent with the cell proportions identified by scRNA-Seq, PVL cells were highly abundant in P4, and lowly detected in P3 (Fig. 2B). PVL cells were highly dispersed throughout the tumour stroma with no obvious co-localisation to the invasive malignant borders. Importantly, our findings suggest that these smooth muscle-like cells, like CAFs, can be readily identified disseminated throughout the stroma, independent of blood vessels.

To understand how the four stromal subpopulations correspond to their normal tissue counterparts, we repeated the staining of PDGFRβ, CD34, α-SMA and CD146 on healthy breast tissue collected from four women. This revealed a high abundance of iCAF-like fibroblasts (PDGFRβ^+^, α-SMA^−^, CD34^HIGH^ and CD146^−^) surrounding ductal regions, while myCAF-like fibroblasts (PDGFRβ^+^, α-SMA^HIGH^ and CD146^−^) were sparsely detected across all four cases (Fig. EV4D). While this small panel of markers do not highlight the large transcriptional changes that may occur upon CAF activation, it does suggest that the broad iCAF-like and myCAF-like fibroblast subsets are resident cell types which are reactivated during carcinogenesis. For PVL cells, IHC staining of CD146 was completely restricted to blood vessels (Fig. EV4D). This further confirmed using co-IF staining for CD31 and CD146 on the normal tissue cases, where CD146 was completely restricted to CD31-positive blood vessels (Fig. EV4E). Our findings suggest that disseminated PVL cells are a distinct feature in a subset of TNBCs.

### Distinct ligand receptor expression predicts diverse stromal crosstalk to the tumour microenvironment

We next sought to investigate how spatially distinct stromal subclasses may interact with other cells within the TME. Here, we annotated our scRNA-Seq dataset using a published set of curated human ligand-receptor pairs [51]. We used these annotations to construct a cell-to-cell communication network and predict intratumoral signalling between the four stromal clusters, and the surrounding neoplastic, immune and endothelial microenvironment. This revealed diverse stromal signalling profiles (Fig. 5A), with myCAFs and iCAFs having the highest overall predicted ligand activity out of all the cell types (Fig. 5B). The ‘interaction strength’, or the weight of each edge, was defined as the product of expression levels of the corresponding ligand and receptor. All ligand-receptor pairs with an arbitrary ‘interaction strength’ cut-off of 0.1 were classified as candidate signalling molecules, which revealed a total of 570, 482, 437 and 357 unique predicted interactions between stromal clusters with cancer epithelial cells, endothelial cells, myeloid cells (Appendix Fig. S3A-C) and T-cell subpopulations, respectively (Appendix Fig. S3D; Appendix Table S3).

**Figure 5.**
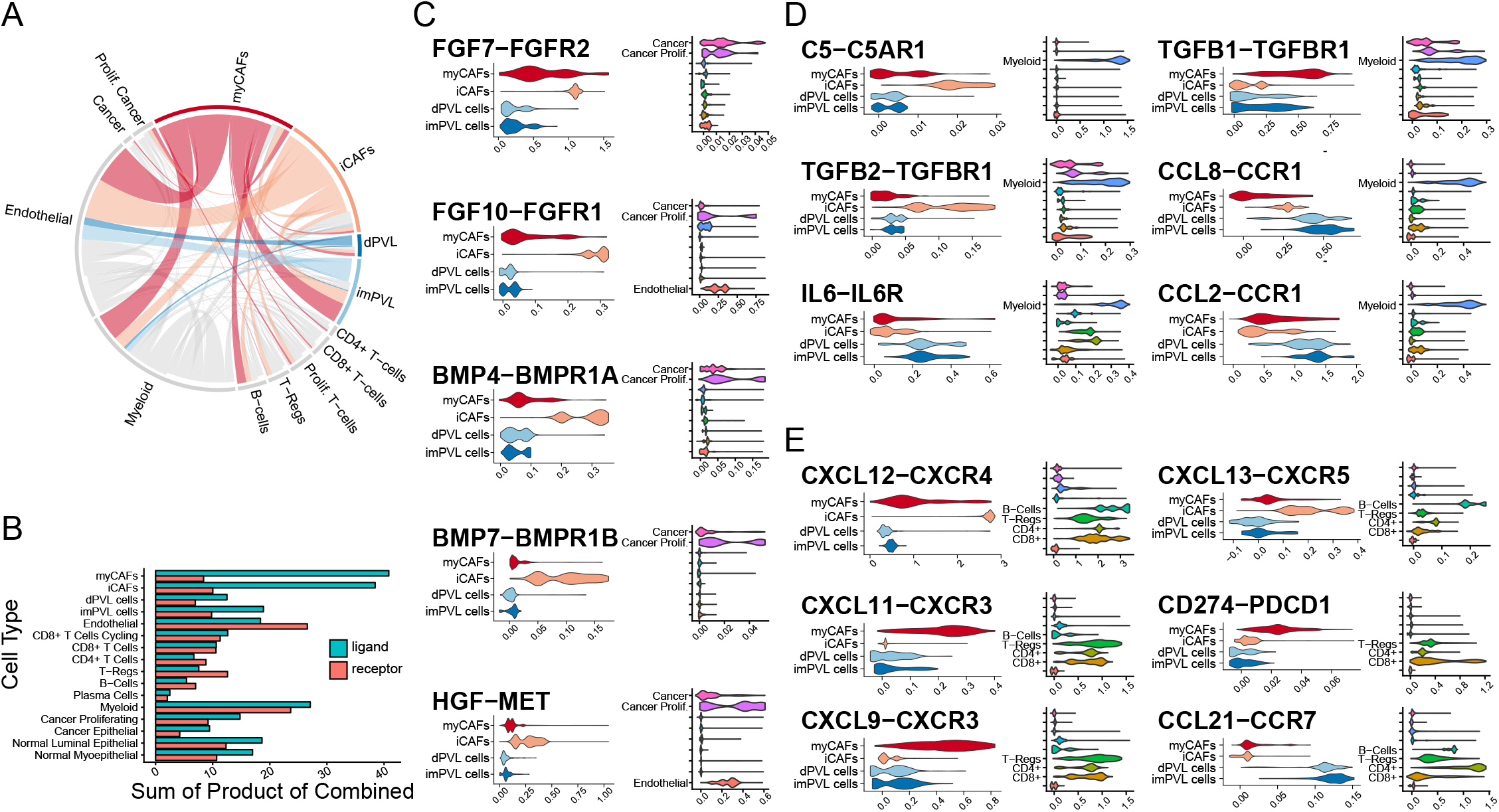
Predicted stromal crosstalk to cancer and immune cells. Overview of the predicted stromal paracrine signalling conserved across the five TNBC patients. The scRNA-Seq dataset were annotated by ligand-receptor pairs as curated in Ramilowski et al. (2015). **a,** Circos plot summary of the stromal ligand-receptor interactions. Outer sectors are weighted according to the number of annotated ligand receptor interactions per cell type. Links between sectors are weighted according to the ‘Interaction Strength’, calculated as a product of ligand and receptor expression. Links are coloured by the respective stromal subsets; myCAFs (red), iCAFs (orange), dPVL cells (blue) and imPVL cells (light blue) **b,** Summary of the total ligands and receptors annotated per cell type. **c-e,** Imputed gene expression of selected candidate signalling molecules identified between the four stromal subsets and malignant **(c)** epithelial, **(d)** myeloid and **(e)** T-cells.

Consistent with the enrichment of growth factor signalling gene ontologies in iCAFs (Fig. 2E), we identified a strong upregulation of crosstalk *via* the FGF (*FGF7* and *FGF10*), BMP (*BMP4* and *BMP7*), HGF and IGF1 pathways to their cognate receptors across cancer cells and endothelial cells (Fig. 5C; Appendix Fig. S3A-B). These factors are known to be highly expressed in breast tumours and associated with breast cancer proliferation, invasion and inducing cancer stem-cell (CSC) phenotypes [52–55]. Different ligands from these pathways were also identified from myCAFs and dPVL cells, suggesting that neoplastic phenotypes could also be influenced by different stromal cells (Appendix Fig. S3A). As we identified iCAFs to be located more distal to the invasive tumour interface, we hypothesize that these secreted factors function from a distance. For signalling to the endothelial compartment, iCAFs and PVL cells were both enriched for well-characterised growth factors involved in angiogenesis (Appendix Fig. S3B). Classical angiogenic pathways including VEGFs (*FIGF*, also known as VEGFD), PDGFs (*PDGFC*), IGFs (*IGF1* and *IGF2*) and Notch signalling (*DLK1*) were enriched in signals emanating from iCAFs (Appendix Fig. S3B). These pathways suggest that the inflammatory CAF phenotype is also associated with tumour neovascularisation [56, 57]. In addition, PVL-derived signals were enriched for the canonical *ANGPT1/ ANGPT2-TIE1* pathway, which are known stimuli that can induce the sprouting of new vessels during the formation of new endothelial tubes [58].

Given the reported immunoregulatory properties of mesenchymal cells [4, 5], we next focused on the signalling of stromal cytokines and checkpoint molecules to immune populations. Here, we identified an enriched interaction between iCAFs and myeloid cells *via* the complement cascade activation interaction *C5-C5AR1* (Fig. 5D; Appendix Fig. S3C). C5 activation in the TME acts as a chemotactic factor for the recruitment of immunosuppressive myeloid cells to suppress T-cell activities [59]. In addition, myCAFs and iCAFs were enriched for *TGFB1-TGFBR1* and *TGFB2-TGFBR1* interactions with myeloid cells, respectively (Fig. 5D; Appendix Fig. S3C). As TGFβ-activated myeloid cells have been shown to enhance breast cancer progression and metastasis *in vivo*, it suggests that both CAF subsets could influence myeloid phenotypes [60]. While the *TGFBR1* receptor was predominantly enriched on myeloid clusters, it is worth noting that its expression was also detected by cancer and endothelial clusters (Fig. 5D). Although PVL cells had lower ligand expression profiles compared to CAFs, several immunomodulatory cytokine interactions were predicted between PVL cells and myeloid cells, including an enrichment of the *CCL8-CCR1, IL6-IL6R* and *CCL2-CCR1* pathways (Fig. 5D; Appendix Fig. S3C). CCL2 produced by the microenvironment in other cancers has been shown to be essential for the recruitment of T-Regs and tumour-associated macrophages, supporting an additional role of PVL cells in recruiting immunosuppressive cells [61].

For the signalling to the lymphocyte compartment, iCAFs had a strong upregulation of the chemo-attractant pathways *CXCL12-CXCR4* and *CXCL13-CXCR5* with T- and B-cells (Fig. 5E; Appendix Fig. S3D). CAF derived CXCL12 has been shown to recruit and regulate the activity of CD4+/CD25+ T-Regs in breast cancers, suggesting iCAFs may have a direct role in recruiting immunosuppressive populations [4, 5]. CXCL12 and CXCL13 signalling axes have also been shown to mediate lymphocyte recruitment to tertiary lymphoid structures (TLS) [62]. MyCAFs were also enriched for secreted immunoregulatory molecules and checkpoints including *CXCL9-CXCR3, CXCL11-CXCR3* and *CD274*-*PDCD1* (PDL1-PD1) with T-cells (Fig. 5E; Appendix Fig. S3D). Lastly, only few candidates were identified between PVL cells with T-cells, including the enrichment of *CCL21-CCR7*, which is associated with immune tolerance in favour of tumour progression (Fig. 5E; Appendix Fig. S3D) [63]. It is evident from our signalling predictions that diverse immunoregulatory molecules are expressed in the stroma, highlighting that immune evasion can be regulated by distinct stromal subpopulations in TNBC.

### Inflammatory-CAFs associated with cytotoxic T-lymphocyte dysfunction

To further investigate the influence of stromal subsets on immune evasion, we explored the association between distinct stromal gene signatures and immune content in three large independent TNBC patient cohorts with associated bulk gene expression data (METABRIC, GSE8812 and GSE21653) [64–66]. Using a computational model called Tumour Immune Dysfunction and Exclusion (TIDE), we examined two primary mechanisms of immune evasion. The first examines factors driving the ‘dysfunction’ of cytotoxic T-lymphocytes (CTLs), while the second examines factors preventing the infiltration of CTLs to the tumour, known as ‘exclusion’ (described below) [67]. TIDE first estimates CTL levels in each sample within a bulk-sequencing cohort using the averaged expression of CTL-specific genes (See Methods). Patients are then stratified into high and low CTL groups based on comparisons to the mean CTL level within the cohort. For dysfunction, we then evaluated whether gene signatures from each of the stromal subsets influences the beneficial effect of CTL levels on patient prognosis [67]. This analysis revealed a strong enrichment of genes from the iCAF signature that were significantly associated with CTL dysfunction in all three bulk tumour cohorts (Fig. 6A). In patients with a low iCAF dysfunction signature level, a significant survival benefit was associated with high CTL levels (Fig. 6B; Fig. EV5A). This is consistent with previous clinical observations in TNBCs where lymphocyte infiltration is a robust prognostic factor for improved disease-free survival and overall survival benefit [68]. Remarkably, in patients with a high iCAF dysfunction signature level, CTL levels were not associated with prognosis in any of the three cohorts (Fig. 6B; Fig. EV5A), suggesting a role for stromal iCAFs in driving dysfunctional CTLs in TNBC. Other stromal subset signatures did not show a significant enrichment of prognostic genes in the context of CTL dysfunction.

**Figure 6.**
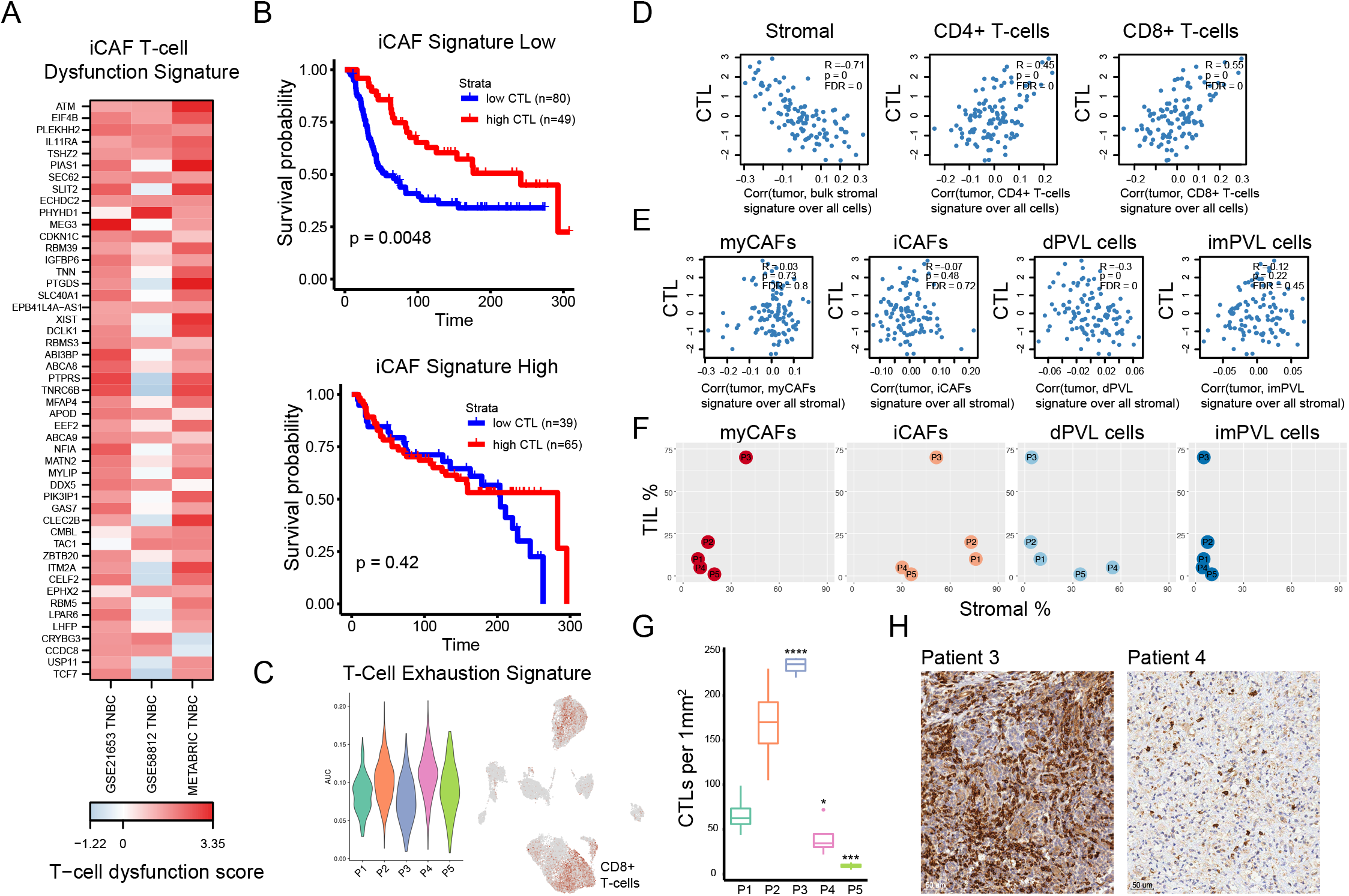
Inflammatory-CAFs and differentiated-PVL cells associated with immune evasion in TNBC patient cohorts. Significant associations between iCAF and dPVL gene signatures with cytotoxic T-lymphocyte (CTL) dysfunction and exclusion in multiple TNBC patient cohorts, respectively, as determined using the tumour immune dysfunction and evasion (TIDE) method. **a,** iCAF T-cell dysfunction gene signature highlighting genes significantly associated with CTL dysfunction in two out of three independent patient cohorts (METABRIC, GSE21653 and GSE58812). **b,** Representative cohort (METABRIC) showing the prognostic value of iCAF T-cell dysfunction signature in the context of CTLs for a total of 233 patients. Kaplan-Meier’s present two groups of patients, ‘low CTL’ (blue line) and ‘high CTL’ (red line), as estimated according to the average expression of CTL-specific genes and stratified as compared to the mean. Tumours with low iCAF T-cell dysfunction signatures (top) show patients with high CTL levels have a better survival outcome. In contrast, this survival benefit is lost in tumours with a high iCAF T-cell dysfunction signature (bottom) **c,** Dysfunctional CTLs detected in all five TNBC patients determined through scoring a T-cell exhaustion signature. UMAP featureplot of the exhaustion signature across all stromal and immune cells as in Fig. 1D. **d,** Bulk stromal signature associates with CTL exclusion. Pearson correlation was computed between all inferred CTL levels (y axis) and the respective correlation between the bulk sample and the single-cell cluster (x axis). Signature of all stromal cells divided over all cells correlated negatively with CTL levels, while control CD4+ and CD8+ gene signatures show a positive correlation. Benjamini-Hochberg procedure was used for adjusting p-values. **e,** dPVL cells associated with CTL exclusion. Repeated analysis in the same manner as in **(d)**, instead with myCAF, iCAF, dPVL and imPVL clusters divided over all stromal cells independently, highlighting that CTL exclusion is mainly driven by dPVL cells. Representative cohort GSE58812 is shown. **f-h,** dPVL profiles and CTL exclusion consistent in our study. **f,** Patients with the highest dPVL profiles by scRNA-Seq (P4 and P5) show the lowest Tumour infiltrating lymphocyte (TIL) pathology counts. **g-h,** Accurate quantification of CTLs and representative immunohistochemistry staining for CD8 on matched patient tumour sections. P3 is shown as an example of a low dPVL profile with high CTLs. In contrast, P4 has a high dPVL profile with low CTLs. *n* = 5 stromal 1 mm^2^ regions were counted per tumour. Statistical significance was determined using pairwise comparison with Student’s *t* test.

To investigate whether CTLs in each patient were indeed dysfunctional, we scored a published T-cell exhaustion gene signature in our CD8+ T-cell populations from each patient using an AUC approach (Fig. 6C) [69]. This gene set includes canonical markers of exhausted T-cells including *PDCD1* (PD-1), *LAG3*, *TIGIT* and *CTLA4* [69]. This revealed heterogeneity for exhausted CD8+ T-cell populations in all 5 patients (Fig. 6C), with P2 and P4 having the highest average exhausted gene signature score. In contrast, the exhaustion signature was not enriched in any other cell population with the exception of the myeloid cell cluster (Fig. 6C). Myeloid cells, which can include tumour-associated macrophages and myeloid derived suppressor cells, are known to hold immunosuppressive properties, and can also express inhibitory molecules associated with T-cell suppression [70].

### Differentiated-PVL cells associated with cytotoxic T- lymphocyte exclusion

We next explored whether particular stromal subsets were associated with CTL exclusion, a cold ‘immune-desert’ phenotype with ‘low CTL’ activity. This was examined using the Pearson correlations between all CTL levels and the respective correlation score between the bulk tumour sample and the single-cell cluster of interest. The averaged expression of all genes from the single-cell cluster are referred to as a signature in this section. Previous studies have reported an association between CAFs and CTL exclusion [67]. Consistent with this, the collective bulk signature from all stromal cells correlated negatively with CTL levels in four TNBC patient cohorts (Fig. 6D; Fig. EV5B). As a positive control, CD4+ and CD8+ T-cell signatures from our dataset positively correlated with CTL levels as expected (Fig. 6D; Fig. EV5B). To investigate if this was predominantly driven by one stromal subset, we repeated this analysis with the averaged gene expression of myCAFs, iCAFs, dPVL and imPVL clusters independently (Fig. 6E; Fig. EV5C). This revealed that dPVL cells were the only subset with a significant negative correlation with CTL level in three of four cohorts, suggesting they are the primary subset associated with T-cell exclusion (Fig. 6E; Fig. EV5C). To further explore this correlation in our five patients, tumour infiltrating lymphocytes (TILs) and CTLs were scored in matched tumour sections by a specialist breast pathologist. Total TILs were estimated using standard H&E-based assessment (Fig. 6F), whilst stromal CTLs were accurately quantified by CD8 staining and scored as previously described (Fig. 6G) [71]. The latter measurements were performed as TILs can also be comprised of non-CTL populations including CD4+ T-cells, T-Regs and B-cells. TILs and CTL scoring revealed that 2 out of 5 patients (P4 and P5) had very low CTL infiltration (<5% TILs and <50 CD8+ T-cells per 1 mm^2^), whereas P3 had a very high infiltration (>70% TILs and >200 CD8+ T-cells per 1 mm^2^) (Fig. 6F-H). In support of dPVL cells as drivers of T-cell exclusion, only 4% of stromal cells from P3 were annotated as dPVL cells, while P4 and P5 had the two largest proportions of dPVL profiles with 35.5% and 26.8% (Fig. 2B). Furthermore, no disseminated PVL cells could be readily detected in P3 using co-IF (Fig. EV4C). While small numbers, our findings are consistent with the proposal that specialised stromal subclasses are associated with immune evasion.

## Discussion

Our study describes a detailed taxonomy of human stromal subclasses in TNBC at cellular resolution. The activated tumour stroma is classically described using a broad ‘CAF’ classification. Here, we provide evidence that it is also comprised of functionally distinct perivascular-like cells which are not necessarily associated with the endothelium. We show that stromal heterogeneity diverges to four distinct states: myofibroblast-like, inflammatory-like CAFs and differentiated- and immature-PVL cells. Similar to CAFs described in pancreatic ductal adenocarcinoma, we find stromal subclasses are spatially distinct, with myCAFs localised to the invasive tumour front, whilst iCAFs are located distal to this interface [12]. From our systematic scRNA-Seq of the TME, we used receptor expression on other cell types to predict diverse stromal-immune crosstalk *via* an array of immunoregulatory molecules to immune populations. We go on to show that iCAF and dPVL subsets are highly associated with immune evasion in multiple independent TNBC cohorts, suggesting a clinical relevance for unique stromal subsets [64–66].

Few studies have investigated the functional heterogeneity of the cancer stroma. A recent scRNA-Seq study profiled CAFs in a mouse model of breast cancer and defined matrix-, vascular-like-, cycling- and developmental-CAF subsets [20]. We did not find a cycling-CAF cluster driven by proliferation markers (Appendix Fig. S4A), likely reflecting unique features of animal models. In addition, the authors proposed mouse ‘developmental-CAFs’ to be of epithelial to mesenchymal transition origin [20]. In contrast, we found the expression of proposed developmental-CAF markers *Scrg1*, *Sox9*, and *Sox10* exclusively in cancer epithelial clusters, which are classified based on the expression of epithelial lineage exclusive keratins (Appendix Fig. S4B-C). Our comparisons suggest that developmental-CAFs are either unique to mouse models or are cancer cells whose expression of *EPCAM* is down-regulated, which was a negative marker used for CAF isolation in these studies [43, 44].

Despite well-characterised roles in cancer progression, the cellular origins of CAFs remain poorly understood. Our results support the notion that dispersed stromal cells can also arise from perivascular cells, likely delaminated from vascular structures. Although PVL cells clustered distinctly from CAFs and express perivascular markers including *MCAM (*CD146*), CAV1, RGS5*, *MYH11* and *TAGLN* (SM-22-Alpha), they also expressed an array of markers commonly used to classify CAFs, including *ACTA2* (α-SMA), *PDGFRB*, *THY1* (CD90), *S100A4* (FSP-1) and *ITGB1* (CD29) [4, 25]. Similar PVL subsets were identified in a previous mouse model of breast cancer [20]. The authors defined these cells as ‘vascular-like CAFs’ through the expression of vessel development markers such as CD146 [20]. Although the authors hypothesised that vascular-like CAFs are derived from perivascular cells such as pericytes, the concept of pericyte-to-fibroblast transition has been debated [72]. Our findings from functional assays suggest PVL cells do not possess the defining fibroblast trait of collagen deposition and remain phenotypically distinct from the fibroblast lineage (Fig. 4E).

The functional role of perivascular cells in breast cancer is poorly understood. A very early study found that 4 out of 10 breast tumours showed substantial infiltration of vascular smooth muscle cells based on staining for markers including α-SMA, smooth muscle myosin and calponin [10]. This finding went without further exploration until this manuscript, where we validate their existence using state-of-the-art scRNA-Seq and staining of CD146 in matched patient tissue. From our TF analysis, we predict the MEF2 regulon to be a strong activating TF of the PVL subclass. MEF2C is a well-defined regulator for establishing vSMCs during development, highlighting a likely vSMC origin of the described PVL cells [39, 40]. As observed during wound healing, we hypothesise that vSMCs could be stimulated by malignant factors or mitogens, allowing them to migrate from the vessel basement membrane into the stroma [73, 74]. This is further supported by *in vitro* studies showing that breast cancer derived PDGFs can induce the recruitment and migration of vSMCs [73]. As perivascular cells play an important part during angiogenesis and blood vessel stability, it is also possible that their displacement in tumours is stimulated by, or a driver of, dysregulated angiogenesis or hypoxia. Although it is yet to be studied in the context of perivascular cells, studies have reported that the imPVL marker CD36 is enriched in normal tissue regions and is associated with good survival outcome in breast cancer [75]. However, the origin and functional role of PVL subpopulations remain to be defined by future studies. The staining of CD146 exclusively associated with blood vessels of normal breast tissue suggests that detached PVL cells are a distinct feature of breast cancers.

Importantly, our findings suggest that previous studies characterising CAFs with a small number of markers have likely also studied PVL cells. For example, subsets discriminated by CD146 have been characterised in endocrine-resistant breast cancers [2]. Patients with a CD146+ stroma demonstrated good responses to tamoxifen therapy through the maintenance of estrogen receptor (ER) dependent proliferation in cancer cells. Our findings suggest that PVL cells rather than CAFs are a biomarker for ER-directed therapeutic response in ER positive breast cancers [2], a prediction that requires more detailed validation. Another elegant study reported a subset of chemoresistance-promoting CAFs, marked by α-SMA+, GPR77+ and CD10+ expression [6]. Due to the shared expression of α-SMA between myCAFs and PVL cells, our findings also raise the question whether PVL cells could also contribute to chemoresistance in a subset of patients [6]. Although we did not find an enrichment of GPR77+ CD10+ α-SMA+ cells in any CAF subclasses, this may be explained by the treatment status of our samples.

Lastly, we found a strong enrichment of immunomodulatory pathways in the predicted signaling between stromal cells and immune cells. We identified an array of important candidates in patient tissue for future experimental studies for functional relevance. It is important to acknowledge, however, that transcript signaling predictions are not always concordant with protein expression. Although no CAF subsets in previous mouse studies were distinguishable by immunomodulatory properties [20], there are several reports of predicted CAF-immune interactions in human tissue. We found that iCAFs expressed an array of immunomodulatory molecules to cognate receptors on T-cells. In other studies, CAFs have been implicated in the recruitment and activity of T-Regs through the regulatory molecules CXCL12, CD40, B7H3, DPP4 and CD73 [4]. In addition, iCAFs also expressed several molecules known to regulate myeloid cells, including complement C5, IL6 and TGFβ [59, 60]. Myeloid cells, including tumour associated macrophages and myeloid derived suppressor cells, are well characterised in contributing to an immunosuppressive TME. Most importantly, gene signatures generated from iCAFs were strongly associated with CTL dysfunction in TNBC patient cohorts. We also report a novel dPVL stromal subset strongly associated with CTL exclusion. We identified an enrichment of dysfunctional/exhausted T-cells which correlated with their respective stromal profiles, though we acknowledge that our study consists of small patient numbers. In patients with the highest dPVL profile, we found consistently low TIL and CD8 counts in matched pathology. Considering the proposed origin of detached PVL cells from the vascular structure, we hypothesise that this may be related to reduced lymphocyte extravasation from dysregulated tumour blood vessels. In support of this, previous studies restoring vascular integrity in tumours through vessel normalisation and increased perivascular coverage find an influx of CD8+ T-cells in tumour tissue [25, 76]. In addition, signalling between CD4+ T-cells and pericytes have also been reported to play a reciprocal role in tumour vessel normalisation [77]. It is possible that the association between dPVL cells and CTL exclusion in patient cohorts reflect tumours with low vascular integrity, and may act as a biomarker for patients suitable for vessel normalisation therapeutic strategies.

Whilst our findings point to the targeting of stromal cells, future work investigating the transcriptional changes in stromal cells between healthy breast tissue and cancer is required to understand the stromal states that are cancer-specific versus reactivated resident cell types. In support of the latter possibility, a recent study showed that there are minimal proteomic differences between normal fibroblasts and CAFs in prostate cancer models [78]. We find that iCAF- and myCAF-like fibroblasts exist in cancer-free normal breast tissue. It is important to note that desmoplasia is often observed in cancer-free tissues, particularly in high risk women with high mammographic density [79]. This can be influenced by several physiological factors such as weight, pregnancy and menopausal status, highlighting important factors that need to be considered in future projects examining the normal breast tissue microenvironment such as the human cell atlas project [79]. These differences may exist from distinct epigenetic states between CAFs and normal fibroblasts, indicating another layer of complexity that remains to be explored in the four breast cancer stromal subsets identified in our study [80]. The integration of future assays combining scRNA-Seq with chromatin states will be important in elucidating the epigenetic regulation of cancer-associated stromal cells. Identifying specific activation markers in comparison to healthy tissue is also an important prerequisite for the development of precise cancer therapeutic strategies with low toxicities. Future *in vitro* and *in vivo* studies will be important in understanding how stromal cells are dynamically reprogrammed and how the subclasses described here restrain or promote tumour growth and invasion.

Clinical trials for mainstream immune checkpoint therapies including anti-PDL1 have shown limited efficacy in the treatment of advanced TNBC. This hints at alternate mechanisms of immune evasion and novel therapeutic strategies are desperately needed to improve immunotherapy for TNBC. Our findings suggest that co-targeting stromal subpopulations could elicit a more effective immune response in a subset of patients through inhibiting CTL dysfunction and exclusion. This remains to be experimentally tested. In conclusion, we have comprehensively profiled four functionally distinct stromal subclasses in human TNBC, not previously described in breast cancer, mouse models or other cancer types. Importantly, we described subsets of CAFs and PVL cells with clinical relevance, presenting as candidates to further investigate. While our dataset captures a majority of the expected cell types from the TME, certain cell types such as adipocytes are under-represented due to biases from standard tissue dissociation protocols. Integration of alternative methods such as single-nuclei sequencing and spatial transcriptomics in future cancer cell atlas studies will be crucial for a comprehensive understanding of the TME. Our findings in only five patients also highlight the potential of applying scRNA-Seq methods to larger scale patient cohorts for the identification of new disease relevant cell states and their gene expression features.

## List of abbreviations

TNBC: Triple-negative breast cancer
CAFs: Cancer associated fibroblasts
PVL: Perivascular-like
scRNA-Seq: Single-cell RNA Sequencing
CTLs: Cytotoxic T-lymphocytes
vSMCs: Vascular Smooth Muscle Cells

## Materials and Methods

### Ethics approval and consent for publication

Patient tissues used in this work were collected under protocols x13-0133, x16-018 and x17-155. HREC approval was obtained through the SLHD (Sydney Local Health District) Ethics Committee; RPAH (Royal Prince Alfred Hospital) zone, and the St Vincent’s hospital Ethics Committee. Site-specific approvals were obtained for all additional sites. Written consent was obtained from all patients prior to collection of tissue and clinical data stored in a de-identified manner, following pre-approved protocols. Consent into the study included the agreement to the use of all patient tissue and data for publication.

### Tissue dissociation

Fresh surgically resected tissue was washed with RPMI 1640 (ThermoFisher Scentific) and minced with scissors. Samples were enzymatically dissociated using Human Tumor Dissociation Kit (Miltenyi Biotec) according to manufacturer’s protocol (https://www.miltenyibiotec.com/AU-en/products/macs-sample-preparation/tissue-dissociation-kits/tumor-dissociation-kit-human.html#gref). Following incubation, the sample was then resuspended in RPMI 1640 and filtered through MACS® SmartStrainers (70 μM; Miltenyi Biotec) and the resulting single cell suspension was centrifuged at 300 × g for 5 min. Red blood cells were lysed with Lysing Buffer (Becton Dickinson) for 5 mins and the resulting suspension was centrifuged at 300 × g for 5 min. Viability was assessed to be > 80% using Trypan Blue (ThermoFisher). Viability enrichment was performed using the EasySep Dead Cell Removal (Annexin V) Kit (StemCell Technologies) as per manufacturers protocol. Dissociated cells were resuspended in a final solution of PBS with 10% fetal calf serum solution prior to loading on the 10X Chromium platform.

### Single-cell RNA sequencing on the 10X Chromium platform

High throughput droplet based SCRS was performed on the single-cell suspensions using the Chromium Single Cell 3’ v2 Library, Gel Bead and Multiplex Kit and Chip Kit (10X Genomics) according the to manufacturer’s instructions, with a target of 5,000 cells per lane. SCRS libraries were sequenced on the Illumina NextSeq 500 platform with pair-end sequencing and dual indexing according to the recommended Chromium platform protocol; 26 cycles for Read 1, 8 cycles for i7 index and 98 cycles for Read 2.

### Data processing

Sample demultiplexing, reference mapping, barcode processing and gene counting was performed using the Cell Ranger Single Cell Software v2.0 (10X Genomics). Reads were aligned to the GRCh38 human reference genome. Raw count matrices were exported and filtered using the EmptyDrops package in R [81]. EmptyDrops distinguishes ‘real’ barcodes from ‘noise’ by calculating deviations of each cell against a generated ambient background RNA profile. Filtered barcodes were then processed using the Seurat v2.0 package in R [21]. Additional conservative cut offs were further applied based on the number of genes detected per cell (greater than 200) and the percentage of mitochondrial unique molecular identifier (UMI) counts (less than 10%). Individual Seurat objects were then integrated using the canonical correlation analysis (CCA) function *RunMultiCCA* according the developer guidelines [82]. The top 2000 most variable genes from each sample were combined for CCA vector identification. The first 20 CC dimensions were used for the alignment of subspaces and UMAP projection.

### Cluster annotation

Cell clusters were annotated using canonical cell type markers for epithelial (*EPCAM*), myoepithelial (*EPCAM*^LO^, *ACTA2, KRT5* and *KRT14*), basal (*KRT5* and *KRT14*), mature luminal (*ESR1*), endothelial (*PECAM1*), immune (*CD45*), T-cells (*CD3D*, *CD8A* and CD4), T-regulatory cells (*FOXP3*), B-cells (*MS4A1*), plasmablasts (*JCHAIN*), myeloid cells (*CD68*) and stromal cells (*PDGFRB* and *COL1A1*). Malignant epithelial cells were distinguished from entrapped normal epithelial cells by inferring copy number variations using the inferCNV package as previously described [22]. In addition, an area under the curve (AUC) approach using published cell type signatures from the XCELL database was performed using AUCell [23, 24]. AUCell scores single cells with input gene signatures and analyses its activity and distribution across the entire dataset to explore the relative expression of the gene set of interest. AUCell utilises raw gene counts and thus, is independent of normalisation bias. CAFs, PVL cells and T-cells were independently re-clustered using the Seurat v3 method. Re-clustering was performed across resolutions 0.2, 0.3, 0.4 and 0.5 to identify stable clusters.

### Differential gene expression and pathway enrichment

The MAST method was used to perform differential gene expression through the *FindAllMarkers* function in Seurat (log fold change threshold of 0.1, *p-value* threshold of 1×10^−5^ and FDR threshold of 0.05). The top 250 DEGs from each cluster were then passed on to the ClusterProfiler package for functional enrichment [32]. The *compareCluster* function was used with the enrichGO databases CC, MF and BP sub-ontologies using the human org.Hs.eg.db database.

### Pseudotime cell trajectory analysis

The Monocle 2 method was applied to infer cell trajectories for CAFs and PVL cells using default parameters, as recommended by developers’ [33]. CAFs from Patient-2 and PVL cells from Patient-1 were extracted for Monocle analysis due to adequate cell numbers and representations of each respective subset. Gene expression matrices from each cell type were first exported from Seurat into Monocle 2 to construct a CellDataSet. Variable genes defined by the differentialGeneTest function (q-val cutoff < 0.001) were used for cell ordering and dimensionality reduction with the setOrderingFilter and reduceDimension functions, respectively.

### Gene-regulatory analysis using SCENIC

Investigation of gene-regulatory networks using SCENIC was performed using a faster python implementation of the tool (pySCENIC) as described by the developers on the 1,729 stromal cells [23, 34]. SCENIC explores gene-regulatory networks by identifying TF co-expression modules and binding motif enrichment. The normalised expression matrix generated from Seurat was first filtered for genes as previously described (sum of gene expression > 3 × 0.005 × 1,729) [18]. Genes detected in at least 0.5% of cells were kept. This resulted in 12,100 genes for pySCENIC input [18]. Analysis was performed using the hg38 mc9nr motif collection with a TSS +/− 10kB (hg38_refseq-r80_10kb_up_and_down_tss.mc9nr) for the arboreto and RcisTarget steps. Gene regulons were clustered and plotted using the pheatmap function in R.

### Flow cytometry and FACS isolation of stromal cells

Cell sorting and flow cytometry experiments were performed at the Garvan-Weizmann Centre for Cellular Genomics, Garvan Institute of Medical Research. Flow cytometry was performed on a Becton Dickinson CantoII or LSRII SORP flow cytometer using BD FACSDIVA software, and the results were analyzed using FlowJo software (Tree Star Inc.). FACS experiments were performed on a FACS AriaIII sorter using the BD FACSorter software. All antibody details used in this study can be found in Supplementary Table S4. Cryopreserved single-cell suspensions from Patient-4 were thawed, washed with RPMI and incubated with an anti-CD16/CD32 antibody (1:200, BD Biosciences #564220) in FACS buffer (PBS containing salts, 2% FBS) for 10 mins to block nonspecific antibody binding. For the isolation of the different stromal subpopulations for subsequent experiments, cells were pelleted and resuspended in FACS buffer containing the following antibodies: anti-EPCAM (1:100; BioLegend #324203), anti-CD31 (1:100; BioLegend #303103), anti-CD45 (1:100; BioLegend #304005), anti-PDGFRβ (1:100; BioLegend #323605), anti-PDGFRα (1:100; BioLegend #323507) and anti-CD146 (1:100; BioLegend #342011) for 20 min on ice. All epithelial, immune and endothelial cells were excluded together on the FITC channel marking EPCAM, CD45 and CD31. In addition, we performed positive selection using PDGFRβ. CAFs and PVL cells were discriminated using PDGFRα and CD146, respectively. CAFs and PVL cells were isolated and cultured into dishes (Corning^®^ LifeSciences) coated with collagen (0.15 mg/ml) in RPMI 1640 supplemented with 20% (v/v) FBS, 50 μg/mL gentamycin and 1x antibiotic/antimycotic (15-240-096, Gibco^®^) in a 5% O_2_, 5% CO_2_ incubator at 37°C. Cell sorting was repeated on cultured CAFs and PVL cells using the previously described experimental conditions with anti-PDGFRα (1:100; BioLegend #323507), anti-CD146 (1:100; BioLegend #342011), anti-FAP (1:100; R&D Systems #FAB3715P-025) and anti-CD36 (1:100; BioLegend #336221). FAP^HIGH^ expression was used to discriminate myCAFs from FAP^LOW^ iCAFs, whilst CD36 expression was used to identify imPVL cells from dPVL cells.

### Immunofluorescence

Primary cells were grown on glass coverslips coated with collagen in the same manner as the CDMs as previously below. Media was removed and cells were rinsed with PBS for 5 min. Cells were fixed in 4% paraformaldehyde (ProSciTech) diluted in PBS for 15 min at room temperature then washed three times with PBS for 5 min. Cells were permeabilised with ice cold methanol for 10 minutes at −20°C followed by three 5 min PBS washes. Cells were blocked in blocking buffer (3% BSA + 0.1% Tween-20 in PBS) for 1 hr at room temperature. Primary antibody was diluted in blocking buffer at the following dilutions: anti-CD34 (1:100; Abcam #MA1-10202), anti-FAPα (1:200; Abcam #ab53066), anti-αSMA (1:500; Abcam #ab21027), anti-CD36 (1:100; Biolegend #336203), anti-CD146 (1:200; Abcam #ab75769), anti-PDGFRβ (1:250; Abcam #ab32570). Coverslips were inverted and incubated on droplets of diluted primary antibody on parafilm in a humidified chamber overnight at 4°C. The following day cells were washed 3 times for 5 min in PBS. Cells were incubated with fluorescent secondary antibody (Jackson ImmunoResearch) diluted 1:500 in blocking buffer for 1 hr at room temperature in a light proof container then washed 2 times with PBS for 5 min. Nuclei were stained with 1 μg/mL Hoechst 33342 (Sigma) in PBS for 5 min at room temperature followed by two 2 min PBS rinses. Coverslips were mounted with Prolong Diamond antifade mountant (Thermo Fisher Scientific) and allowed to dry overnight at room temperature. Fluorescent images were captured using a Leica DMI Sp8 confocal microscope.

Immunofluorescence was performed on 4 μm FFPE tissue sections prepared as described below for IHC. Antigen retrieval was performed for 20 min in a 100°C water bath in target retrieval buffer, pH9 (Agilent Technologies). Slides were blocked for 1 hr at room temperature in PBS containing 3% BSA and 5% goat serum. Slides were incubated with primary antibodies diluted in blocking buffer: anti-CD31 (1:50; Agilent Technologies #M0823) and anti-CD146 (1:600; Abcam #ab75769). Secondary antibody staining, nuclear counterstaining and microscopy were performed as described above.

### Quantitative real time PCR analysis

RNA was extracted from bulk and sorted CAF cells using the Qiagen miRNeasy mini kit (Qiagen) and was reverse transcribed using the Transcriptor First Strand cDNA synthesis kit (Roche). TaqMan assays (Thermo Fisher Scientific) were used to analyse mRNA expression levels using a QuantStudio 7 Flex RT PCR machine (Thermo Fisher Scientific). TaqMan probes used were *FAP* (Hs00990791_m1), *ACTA2* (Hs00426835_g1), CXCL12 (Hs00171022_m1), *EGFR* (Hs01076078_M1), *PDGFRA* (Hs00998018_m1), *PDGFRB* (Hs01019589_m1) and *ACTB* (Hs99999903_M1). Relative gene expression was calculated using the ΔΔCt method.

### Cell Derived Matrices (CDMs)

CDMs were established as previously described [45]. A total of 1.5×10^5^ cells/well were allowed to expand until confluent and ascorbic acid (50mg/ml) added to culture medium on days one, three and five. To maintain the structure interact of the matrix architecture CDMs, were imaged using Second Harmonic Generation (SHG) at Day seven with cells still present in the matrix.

### Second Harmonic Generation (SHG) imaging

Second Harmonic Generation (SHG) Imaging was achieved using an inverted Leica DMS 6000 SP8 confocal microscope with a Ti-Sapphire femtosecond laser cavity (Coherent Chameleon Ultra II) excitation source, operating at 80 MHz and tuned to a wavelength of 880 nm, as previously described [83–85]. SHG intensity was detected using a 440/20 nm RLD HyD detectors. For CDMs 3 representative fields of view (512 μm × 512 μm) were imaged over a 3D z-stack (80 μm with a 2.52 μm step size, and 30 μm with a 1.51 μm step size, respectively), with a line average of 4 at 25x magnification. Rotation images were acquired on the z-level of maximum intensity with a line average of 64 at 63x magnification.

### Collagen fibre orientation analysis

Collagen fibre orientation analysis in SHG images from plugs was carried out as previously described [3, 86]. Briefly the distribution of orientation of collagen within images was assessed based on methodology published by Rezakhaniha et al [87]. The local orientation and isotropic properties of individual pixels making up collagen fibres were derived from structure tensors evaluated by computing the continuous spatial derivatives in the x and y directions using a cubic B-spline interpolation to obtain the local predominant orientation. Graphical outputs show a hue-saturation-brightness (HSB) color-coded map indicating the angles of the oriented structures within the image. Orientation distribution peaks were then aligned. The shape of the distribution indicates the degree of alignment within the image, where wide and broad shapes suggested little coherency in alignment, and tight peaks with small standard deviations implied aligned structures.

### *Immunohistochemistry* and *image alignment*

In house FFPE blocks were made of patient tissues by fixing in 10% neutral buffered formalin for 24hrs and processing for paraffin embedding. Where tissue was limited, diagnostic tumour FFPE blocks were accessed for analysis. FFPE blocks were sectioned at 4 μm. These were used for histological analysis, using a standard Haematoxylin and Eosin stain, and for immunohistochemical analysis on the Leica BOND RX Autostainer. Details of antibodies and staining conditions are described in Supplementary Table 4. H&E and IHC slides were imaged using the Aperio CS2 Digital Pathology Slide Scanner. IHC images were imported into FIJI as a virtual stack. Each layer was then aligned using least squared mode (linear feature correspondences), propagating to the first and last layers for rigid transformation. All other parameters were set to default in FIJI.

### Cell-signalling predictions using ligand receptor annotation

Genes from the scRNA-Seq data were annotated based on a published set of human ligand-receptor pairs derived from supporting literature [51]. We used this knowledge to construct a cell-to-cell communication network between the four stromal clusters and other epithelial, immune and endothelial clusters. To investigate conserved signalling modules in TNBCs, we applied this to the cluster averaged expression levels of all ligands and receptors in the integrated dataset of five patients. The ‘interaction strength’, or the weight of edges between two clusters, was defined as the product of expression values from the ligand and its cognate receptor. All ‘interaction strengths’ greater than an arbitrary cut-off of 0.1 were considered as cell signalling candidates and kept for subsequent analyses (Table S3). The total number of interaction pairs identified per cluster were used to generate summaries of this data (Fig. 5A-B). The top 100 candidates between the four stromal subsets and each target population were clustered using hierarchical clustering (complete and Euclidean distance) and rescaled for visualisation in ggplot2. For the visualisation purposes only, the ligand and receptor expression values in Figure 5C-E were imputed using the MAGIC method to better represent the structure of genes with low expression and dropout [88]. Raw count matrices and cluster IDs identified by Seurat (as previously described) were used as input to MAGIC and run with default parameters as recommended by the developers.

### T-cell dysfunction and exclusion analysis

To investigate the immunomodulatory roles of different stromal subsets, we performed T-cell dysfunction and exclusion analysis using similar strategy from TIDE [67]. We first used the average expression level of *CD8A*, *CD8B*, *GZMA*, *GZMB* and *PRF1* to estimate the cytotoxic T lymphocyte (CTL) level in each sample from the bulk sequencing cohort. Patients with a higher and lower CTL level compared to the mean CTL level within the cohort were stratified into high and low CTL groups, respectively. For CTL dysfunction analysis, TIDE evaluates whether gene signatures from each of the stromal subsets influences the beneficial effect of CTL levels on patient prognosis. This is performed using the interaction coefficient d from Cox proportional hazard (Cox-PH) model to evaluate how the interaction between a candidate gene and the CTL affects the death hazard. Genes with a higher TIDE dysfunction score suggests antagonistic interactions with regards to CTL levels, where the survival benefit of patients with high CTL is lost and thus, suggesting an association with CTL dysfunction. This method was used to calculate the TIDE T-cell dysfunction score from the differentially expressed genes across the four stromal subsets in TNBC patients from the METARBRIC cohort [64] and two independent TBNC cohorts [65, 66]. A total of 233, 84 and 107 patients were evaluated for the METABRIC, GSE21653 and GSE58812 cohorts, respectively.

For T-cell exclusion analysis, we examined Pearson correlations between all CTL levels (indicated on the y axis in Fig. 6E) and the respective correlation score between the bulk tumour sample and single-cell cluster of interest (indicated on the x axis in Fig. 6E). Here, gene signatures for the single-cell cluster of interest were defined by the averaged gene expression of all single cells in the cluster, divided over the averaged gene expression of all cells detected in the dataset. This method was used to define signatures in this section, as opposed to a DEG list in the previous CTL dysfunction analysis. This was first performed for all stromal cells, CD4+ and CD8+ T-cell clusters divided over all detected cells independently, as shown in Figure 6D. We next repeated this for the myCAF, iCAF, dPVL and imPVL clusters divided over all stromal cells independently, as shown in Fig. 6E. In each of the breast cancer cohorts, a higher correlation suggests a positive association between the single cell cluster of interest and CTL levels (as observed in CD4+ and CD8+ T-cells shown in Fig. 6D), while a negative correlation suggest a negative association (as observed in dPVL cells shown in Fig. 6E). This correlation indicates a potential affluence of each stromal subset on T-cell infiltration in tumours. For T-cell exclusion analysis, we examined the three aforementioned TNBC cohorts, as well as the TNBC cohort from The Cancer Genome Atlas (https://www.cancer.gov/tcga) [89].

## Data availability

The scRNA-Seq data from this study has been deposited in the European Nucleotide Archive (ENA) under the accession code PRJEB35405. This depository includes the demultiplexed paired ended reads (R1 and R2), Illumina indices and bam files processed using the Cellranger software. The scRNA-Seq analysis scripts can be found on the website: https://github.com/sunnyzwu/stromal_subclasses. All relevant data are available from the authors upon request.

## Acknowledgements

This work was supported by funding from John and Deborah McMurtrie, the National Breast Cancer Foundation (NBCF) of Australia; and The Sydney Breast Cancer Foundation. A.S. is the recipient of a Senior Research Fellowship from the National Health and Medical Research Council of Australia. S.Z.W. is supported by the Australian Government Research Training Program Scholarship. S.O.T. is supported by the NBCF (PRAC 16-006), the IIRS 19 084 and the Sydney Breast Cancer Foundation and the Family and Friends of Michael O’Sullivan. T.R.C. is supported by an NHMRC RD Wright Biomedical Career Development Fellowship, a Susan G Komen Career Catalyst Award and a Cancer Institute NSW (CINSW) fellowship. S.J. is supported by a research fellowship from the NBCF. X.S.L. is a supported by the Breast Cancer Research Foundation (BCRF-19-100) and the National Institute of Health of the United States (R01CA234018). We would like to thanks the following people for their assistance in the experimental part of this manuscript; Ms. Gillian Lehrbach from the Garvan Tissue Culture Facility; Ms. Anaiis Zaratzian from the Garvan Histopathology Facility for tissue processing and IHC staining; The Garvan-Weizmann Centre for Cellular Genomics, including Mr. Eric Lam, Ms. Hira Saeed and Ms. Melissa Armstrong for the expertise in flow sorting, and Mr. Dominik Kaczorowski for his help in next-generation sequencing.

## Author Contributions

A.S. conceived the project and directed the study with input from all authors. S.Z.W. and A.S. wrote the manuscript with input from all authors. Clinical collaborators E.L., S.W., M.N.H., C.C., C.M., D.S., E.R., A.P., J.B. and L.G organised the access to patient tissue. S.Z.W., G.A. and K.H. optimized the tumour dissociation for scRNA-Seq. C.C. helped perform the next-generation sequencing of the scRNA-Seq libraries. S.Z.W. interpreted and performed the pre-processing and downstream analysis of the scRNA-Seq data. D.R. supervised the scRNA-Seq analysis. J.T. helped with the inferCNV analysis. N.B. helped with the IHC image analysis and alignment. K.H. helped performed the IHC staining experiments. S.Z.W., A.C., K.H. and G.A. helped perform flow sorting experiments. K.J.M. and B.P. performed the collagen assays and imaging. T.R.C helped analyse the collagen fibre orientation data. H.H. performed the IF experiments. C.W. and X.S.L. performed and helped interpret the T-cell dysfunction and exclusion and patient survival analysis. S.O.T. independently scored the H&E and IHC stains. A.F. and R.H. assisted in the analysis of the cell signalling data. T.R.C and P.T. provided intellectual input and helped with the interpretation of the ECM assay data. E.L., S.J. and J.P. provided intellectual input.

## Conflicts of interests

No competing interests.

## Appendix Table Legends

**Appendix Table S1. Differentially expressed genes across the four stromal subsets.** Performed using the MAST method through the *FindAllMarkers* function in Seurat. A log fold change threshold of 0.1 and a *p-value* threshold of 1×10^−5^ and FDR threshold of 0.05 was used.

**Appendix Table S2. Gene ontology pathways enriched across the four stromal subsets.** Functional enrichement was performed using the ClusterProfiler package with the top 250 differentially expressed genes from each stromal cluster. The *compareCluster* function was used with the enrichGO databases CC, MF and BP sub-ontologies using the human org.Hs.eg.db database.

**Appendix Table S3. Predicted stromal paracrine signalling.** Ligands and receptors as annotated from Ramilowski et al. (2015). The interaction strength was defined as the product of the average log normalised gene expression values ligand and receptor values from each cluster. Interactions were rescaled by the interaction pair.

**Appendix Table S4. Antibodies details.** Details of the commercial antibodies used for FACS, immunofluorescence and immunohistochemistry.

## Expanded View Figure Legends

**Figure Expanded View 1. Clinical pathological features and overview of single cell RNA sequencing metrics. a,** Clinical and pathological features of patient age, breast cancer subtype, tumour grade, Ki67 status, treatment history and TIL count of the 5 primary breast carcinoma samples analysed in the study. **b,** Representative hematoxylin-eosin (H&E) stained sections for each patient analysed by single-cell RNA sequencing in this study. **c,** Quality control metrics as generated by the Cellranger software (10X Genomics). **d,** Number of cells that passed quality control and filtering using EmptyDroplets per patient. **e,** Number of cells that passed quality control and filtering using EmptyDroplets per cell type and patient. **f-h,** Number of detected genes **(f)**, UMIs **(g)** and proportion of mitochondrial counts **(h)** per cell type across all samples, respectively.

**Figure Expanded View 2. Scoring of cell type signatures for cluster annotation and re-clustering of T-cells. a,** Featureplots highlighting the area under the curve (AUC) value for selected cell type signatures derived from various studies collated in the XCELL study. AUC values are calculated on a per cell basis using the AUCell package with default parameters. Selected signatures for fibroblasts (Fantom_1), endothelial cells (Fantom_2), B-cells (Fantom_1), Plasma cells (IRIS_2), CD4+ T cells (Fantom_3), CD8+ T cells (HPCA_3), T-regulatory cells (Fantom_3) and monocytes (Fantom_3). **b-d,** Reclustering of 7,990 T-cells identifies 175 T-follicular helper cells (2.2%; *CXCL13* and *CD200*), 994 T-Regulatory cells (12.4%; *FOXP3* and *BATF*), 2,003 other CD4+ T-cells (25.1% of all T-cells; *CD4, IL7R* and *CD40LG*), 3,691 CD8+ T-cells (46.2%; *CD8A* and *GZMH*), 605 proliferating T-cells (7.6%; *MKI67*), 358 NK Cells (4.5%; *GNLY*, *KLRD1*, *NCR1, XCL1 and NCAM1*) and 164 NKT cells (2.1%; *GNLY, KLRD1, NCR1 and CD3D^−^*). Shown are *t-*SNE representations of reclustered T-cells coloured by the annotated subsets **(b)** and patient ID **(c)**. **d,** Heatmap of the top 10 DEGs per T-cell subset. **e,** AUC values for all stromal cells scored against published human and mouse pancreatic CAF signatures for the myofibroblast-like CAF, inflammatory-like CAF and antigen-presenting CAF subsets [12–14]. **f,** Heatmap of the stromal cluster averaged expression of genes from human pancreatic CAF signatures, as in **(e)**.

**Figure Expanded View 3. TNBC stromal subsets per patient, pseudotime trajectory and validation of stromal cultures. a,** *t-*SNE representation of the four subclasses of cancer-associated fibroblasts (CAFs) and perivascular-like (PVL) cells by patient ID. **b,** Differential gene expression heatmaps showing the composition of the four stromal subsets in Patient-1 to Patient-5. **c-d,** Pseudotime trajectory of CAFs from Patient-2 **(c)** and PVL cells from Patient-1 **(d)** using the Monocle method annotated by the subsets derived from Seurat based re-clustering. **c,** Increased expression of marker genes such as *ACTA2*, *COL1A1*, *FAP*, *TAGLN* and *THY1* as cells move throughout pseudotime indicate that iCAFs transition towards myCAFs. In contrast, iCAF marker *CXCL12* decreases as cells move throughout pseudotime. **d,** Increased expression of marker genes such as *ACTA2 and MYH11* as cells move throughout pseudotime indicate that imPVL cells transition towards dPVL cells. In contrast, imPVL cell markers *CD36* and *RGS5* decreases as cells move throughout pseudotime. **e,** Four technical replicates of CAF sorting of myCAF and iCAF fractions using FAP^HIGH^ and FAP^negative/LOW^, respectively. **f,** FACS analysis showing the co-expression of CD90 (THY1) with FAP^HIGH^ <u>CAFs</u>. This is represented through overlaying CD90 signal over a replicate sample used for FACS as in **(e)** (top) and through a contour plot of FAP vs CD90 signal (bottom). **g,** Quantitative-PCR validation of *FAP, ACTA2, CXCL12, EGFR, PDGFRA* and *PDGFRB* in bulk, FAP^HIGH^ and FAP^negative/LOW^ CAF sorted fractions. Consistent with scRNA-Seq findings, *FAP* and *ACTA2* are enriched in FAP^HIGH^ sorted myCAF-like fractions, while *CXCL12 and EGFR* are enriched in FAP^LOW^ sorted iCAF-like fractions.

**Figure Expanded View 4. Immunohistochemistry and immunofluorescence of human breast cancers and normal breast tissue. a-b,** Additional immunofluorescence images of cultured human CAFs **(a)** and PVL cells **(b)**, staining for CD34 (CAFs), α-SMA (myCAFs and PVL cells), CD146 (PVL cells) and CD36 (imPVL cells). **c,** Validation of PVL cells detached from blood vessels using co-immunofluorescence of CD31 (red), CD146 (green) and DAPI (blue) for sections from all five patients analysed in this study. **d,** Immunohistochemical staining of PDGFRβ, α-SMA, CD34 and CD146 in serial sections cut 4 μm apart from normal breast tissues collected from four women. Images were aligned using FIJI. **e,** Co-immunofluorescence of CD31 (red), CD146 (green) and DAPI (blue) from normal breast tissue samples. CD146 is completely colocalised with CD31, suggestion no detached PVL cells are present in normal breast tissues.

**Figure Expanded View 5. Influence of inflammatory-CAF and differentiated-PVL subclasses on T-cell dysfunction in TNBC patient cohorts.** The association between stromal gene signatures, cytotoxic T-cell levels, and overall patient survival in all three TNBC patient cohorts examined in this study (METABRIC – 233 patients, GSE21653 – 84 patients and GSE58812 – 107 patients). Using the TIDE method, we show significant associations between iCAF and dPVL gene signatures with cytotoxic T-lymphocyte (CTL) dysfunction and exclusion. **a,** Prognostic value of iCAF T-cell dysfunction signature in three independent cohorts. Kaplan-Meier’s present two groups of patients, ‘low CTL’ and ‘high CTL’, as estimated according to the average expression of *CD8A*, *CD8B*, *GZMA*, *GZMB* and *PRF1*, and stratified as compared to the mean. The top and bottom panels show tumours with low and high iCAF T-cell dysfunction signature, respectively. Sample divided according to iCAF T-cell dysfunction signature show significant association with CTL levels and survival outcome. **b,** Bulk stromal signature associates with CTL exclusion. Pearson correlation was computed between all inferred CTL levels (y axis) and the respective correlation between the bulk sample and the single-cell cluster of interest (x axis). Signature of all stromal cells divided over all cells correlated negatively with CTL levels, while control CD4+ and CD8+ gene signatures show a positive correlation. Benjamini-Hochberg procedure was used for adjusting p-values. **c,** dPVL cells associated with CTL exclusion. Repeated analysis in the same manner as in **(b)**, instead with the averaged expression signature of each stromal subset over all stromal cells, highlighting that CTL exclusion is mainly driven by dPVL cells in three out of four cohorts. TNBC data cohort from The Cancer Genome Atlas (TCGA) was also examined for CTL exclusion analysis.

## Appendix Figure Legends

**Appendix Figure S1. Identification of malignant cancer cells using inferCNV. a-d,** Inferred copy number variation profiles as estimated using the inferCNV method. Epithelial cells in each dataset with distinct copy number profiles were classified as cancer for downstream cell-signalling analysis with each stromal subset. Only epithelial cells are highlighted in P1 due to low gene coverage for inferCNV analysis.

**Appendix Figure S2. Top transcriptional activators distinguishing the four stromal subpopulations. a,** The log normalised gene expression (left) and respective AUC regulon activity (right) for the top 50 highest TFs regulons as estimated using SCENIC. Regulons are all filtered for TFs that were statistically enriched between the four subsets (p < 1×10^−5^ One-way ANOVA). **b,** Correlation strengths between the log normalised gene expression and regulon activity (AUC) for the top 50 highest TFs regulons as estimated using SCENIC. Regulons are all filtered for TFs that were statistically enriched between the four subsets (p 1×10^−5^ One-way ANOVA). R-squared values were computed using linear regression in R.

**Appendix Figure S3. Signalling between stromal subsets and cancer epithelial, endothelial, myeloid and T-cell subpopulations.** Hierarchical clustering (Euclidean and complete distance) of the top 100 candidate signalling molecules between the four stromal populations and **(a)** cancer cells and proliferating cancer cells, **(b)** endothelial, **(c)** myeloid cells and **(d)** T-cell subsets including CD8+ T-lymphocytes, cycling T-lymphocytes, CD4+ T-lymphocytes and T-regulatory cells. Ligand and receptor pairs were ranked according to the ‘interaction strength’, defined as the product of ligand and receptor expression. All interaction strength values were rescaled per interaction.

**Appendix Figure S4. Mouse models of breast cancer do not completely recapitulate human stromal subsets. a,** Violin plot highlighting the negative expression of the proliferation markers *MKI67* and *AURKA* in the four CAF and PVL subsets, highlighting that cycling-CAFs may be unique to aggressive mouse models. **b-c,** Log normalised expression of the previously reported mouse developmental CAF markers *SOX9*, *SCRG1* and *SOX10,* and epithelial markers *EPCAM*, myoepithelial markers *KRT5, KRT14* and *ACTA2,* showing exclusive expression in epithelial clusters rather than in stromal populations.

## Notes

### Competing Interest Statement

The authors have declared no competing interest.

