## Supplementary Figures for "Single-cell analysis reveals diverse stromal subsets associated with immune evasion in triple-negative breast cancer"

### Expanded View 1

A

| | Patient 1 | Patient 2 | Patient 3 | Patient 4 | Patient 5 |
|---|---|---|---|---|---|
| Age | 35 | 49 | 47 | 73 | 67 |
| Subtype | TNBC | TNBC | TNBC | TNBC (Metaplastic) | TNBC |
| Grade | III | III | III | III | III |
| Ki67 | 70% | 40% | - | 75% | 60% |
| Treatment | No | No | No | Treated (AC) | No |
| TIL count | 10% | 20% | 70% | 5% | 1% |

B

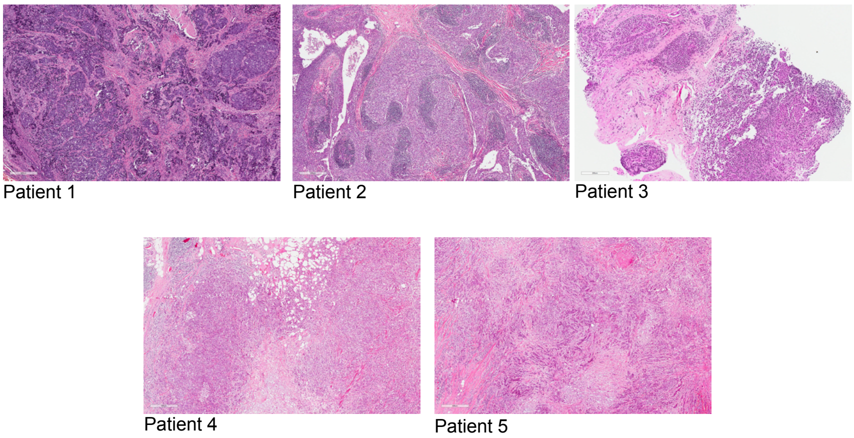


C

| | P1 | P2 | P3 | P4 | P5 |
|---|---|---|---|---|---|
| Number of Reads | 107.8 M | 216.2 M | 152.9 M | 152.8 M | 194.6 M |
| Q30 Barcode (%) | 97.6 | 97.6 | 96.7 | 97 | 96.3 |
| Q30 RNA Read (%) | 83.7 | 86.9 | 86.2 | 88.3 | 68.9 |
| Q30 Sample Index (%) | 95.7 | 97 | 96.5 | 96.8 | 91.6 |
| Q30 UMI (%) | 96.6 | 96.7 | 96 | 96.3 | 96.1 |
| Sequencing Saturation (%) | 82.9 | 43.8 | 25 | 37.2 | 41.9 |
| Total Genes Detected | 20,833 | 23,563 | 23,336 | 22,194 | 22,649 |

D

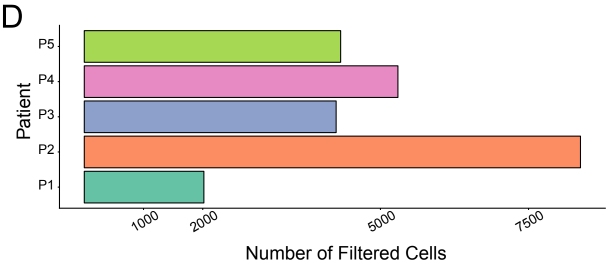


E

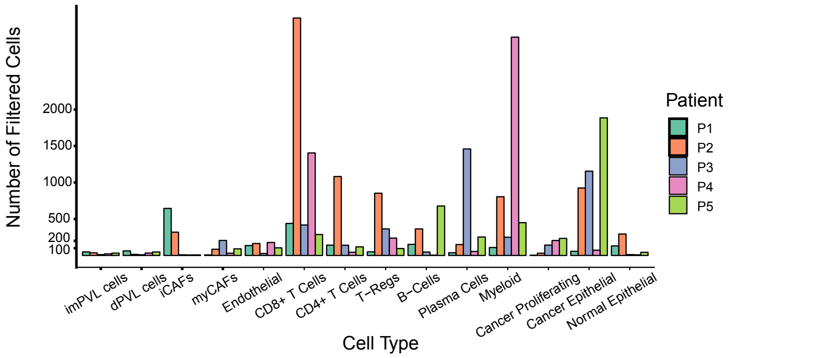


F

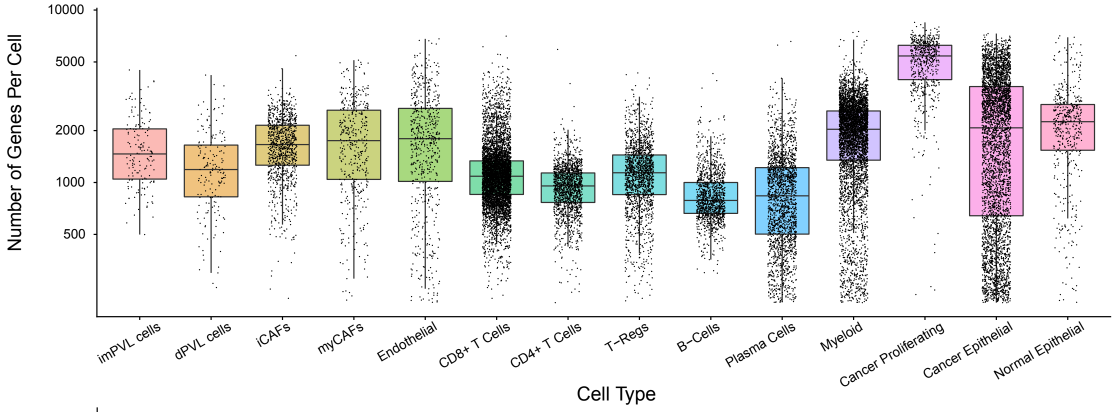


G

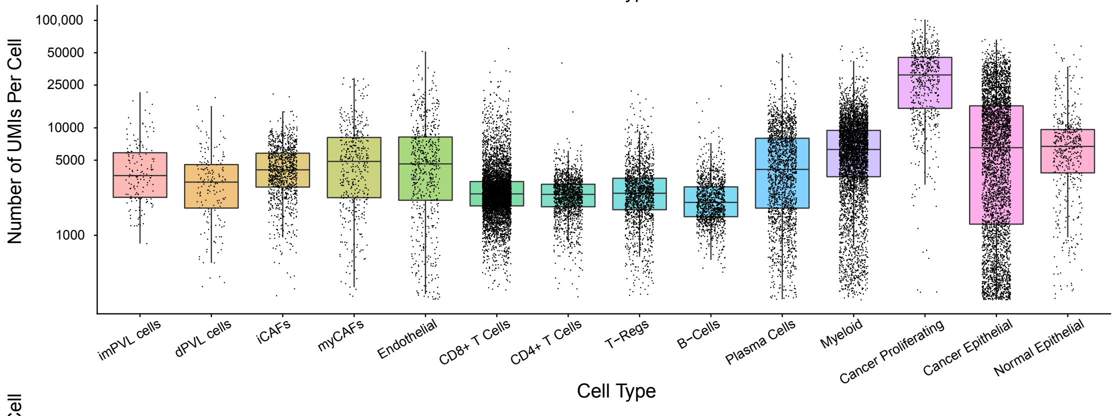


H

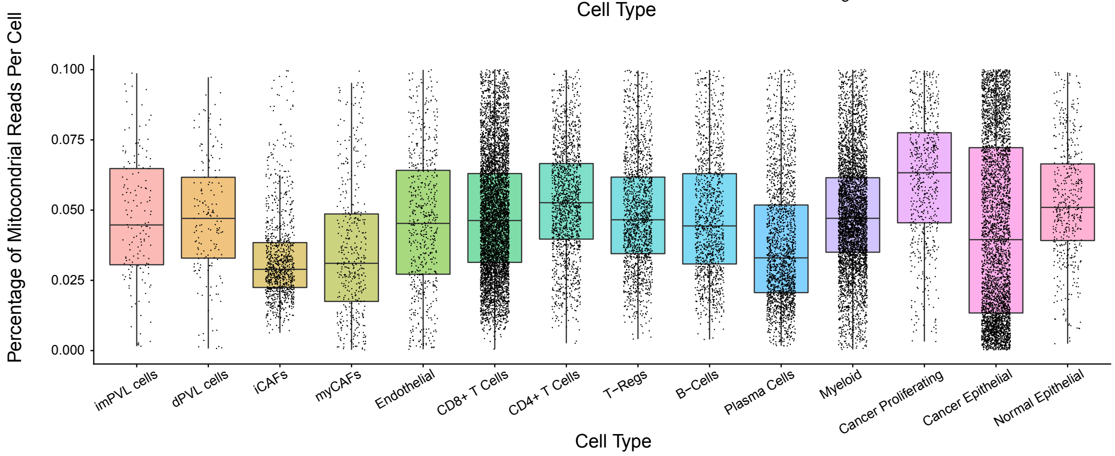

### Expanded View 2

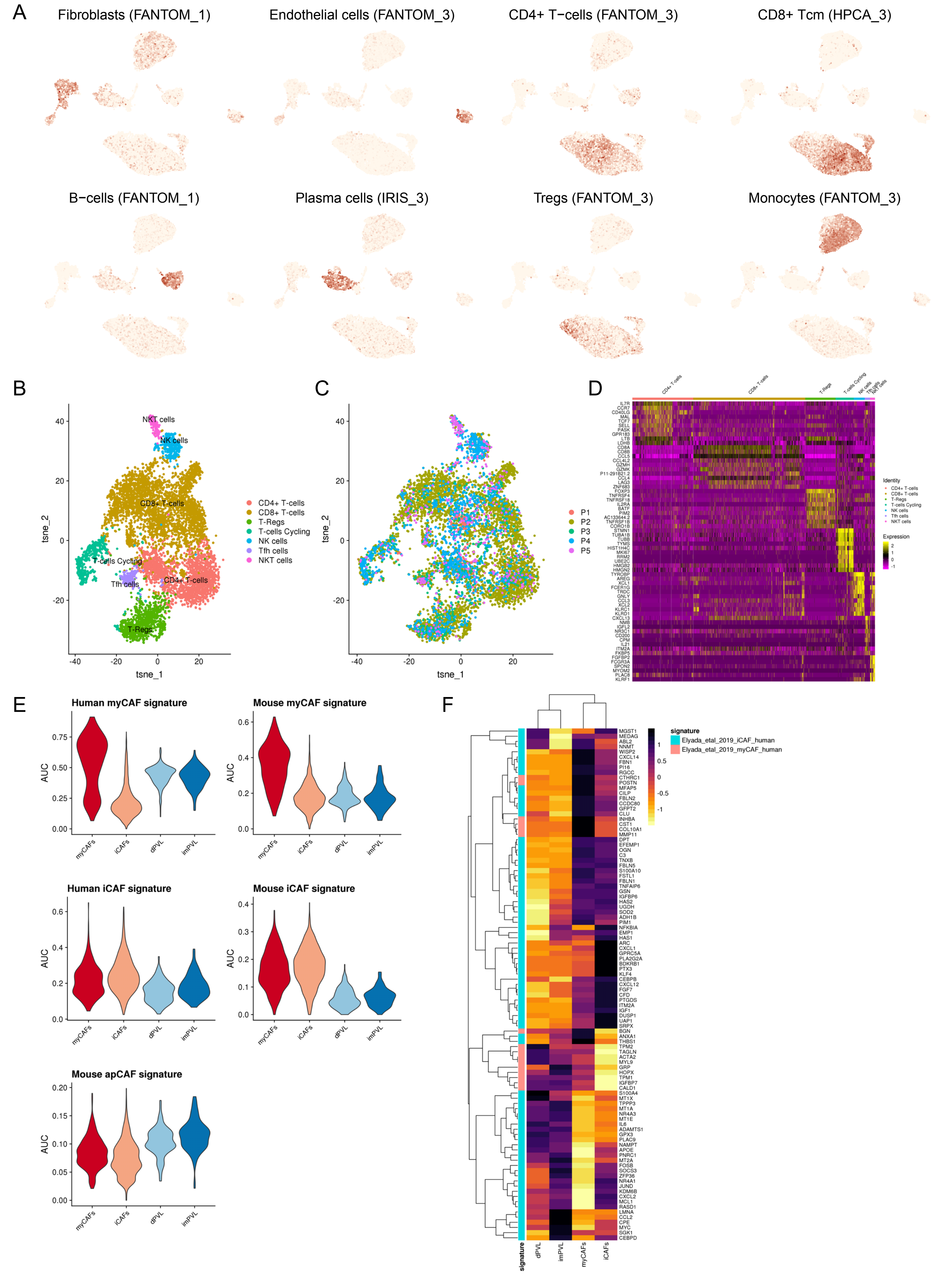

Expanded View 3

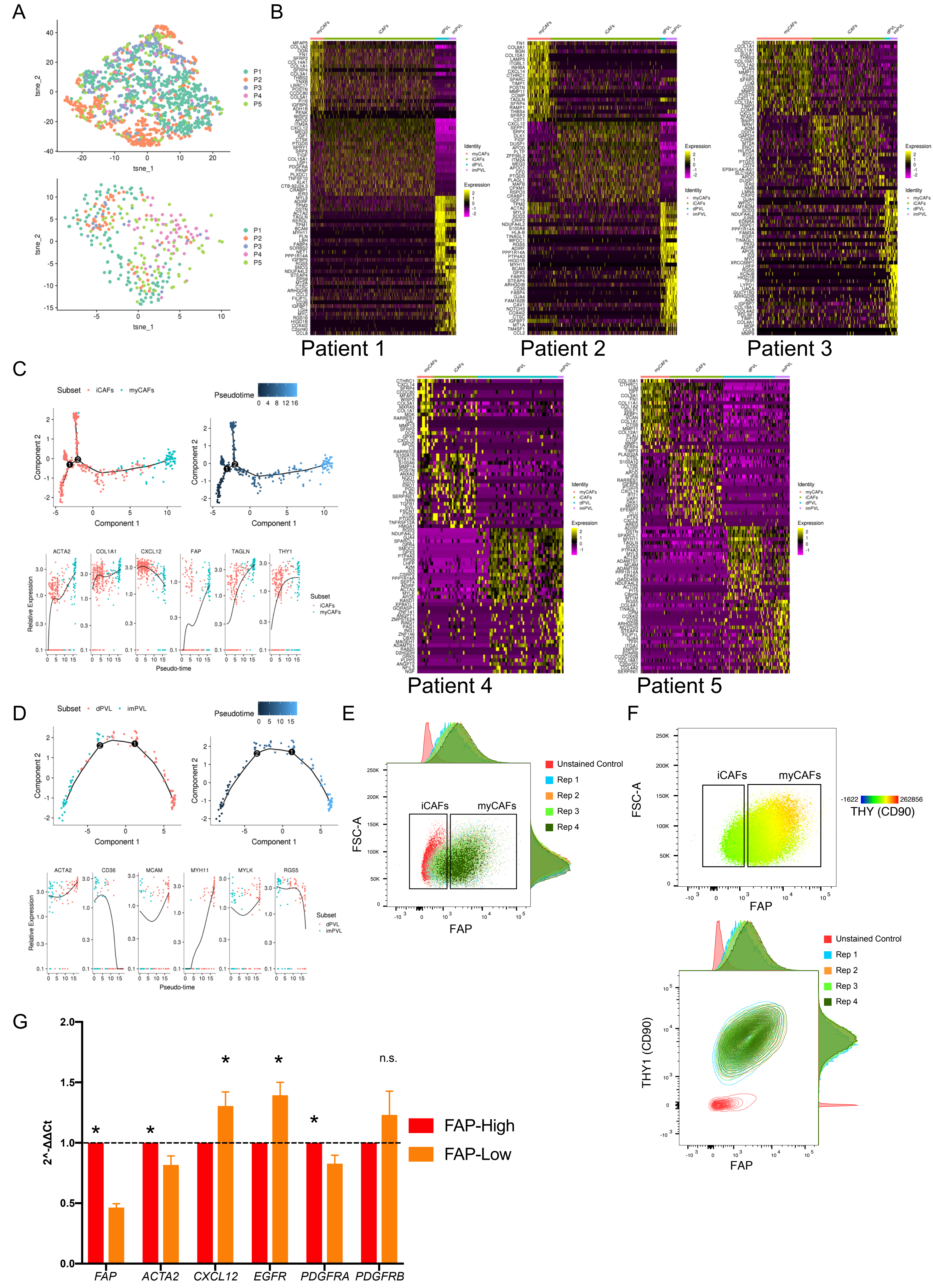

### Expanded View 4

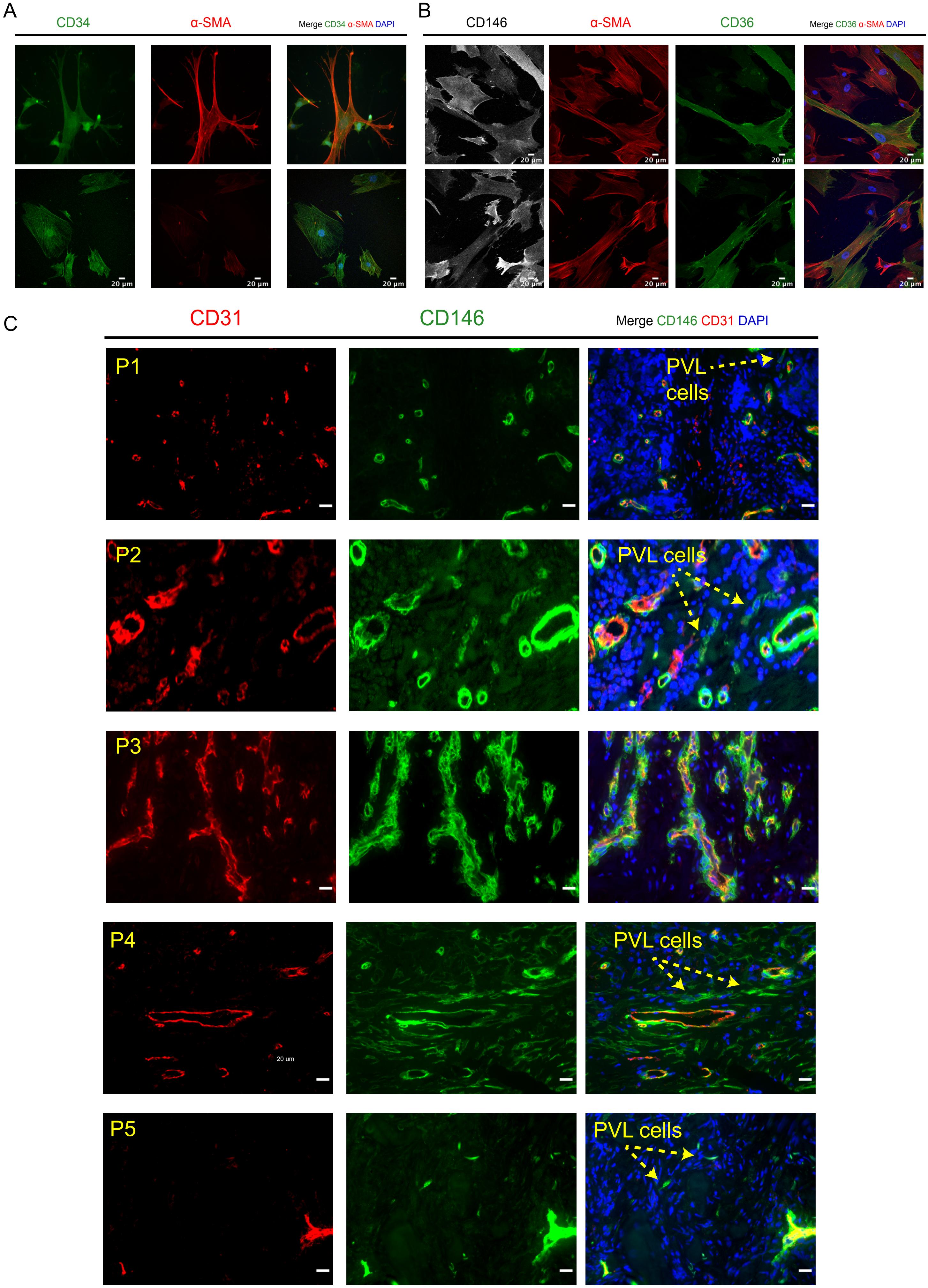

Expanded View 4 (cont.)

D

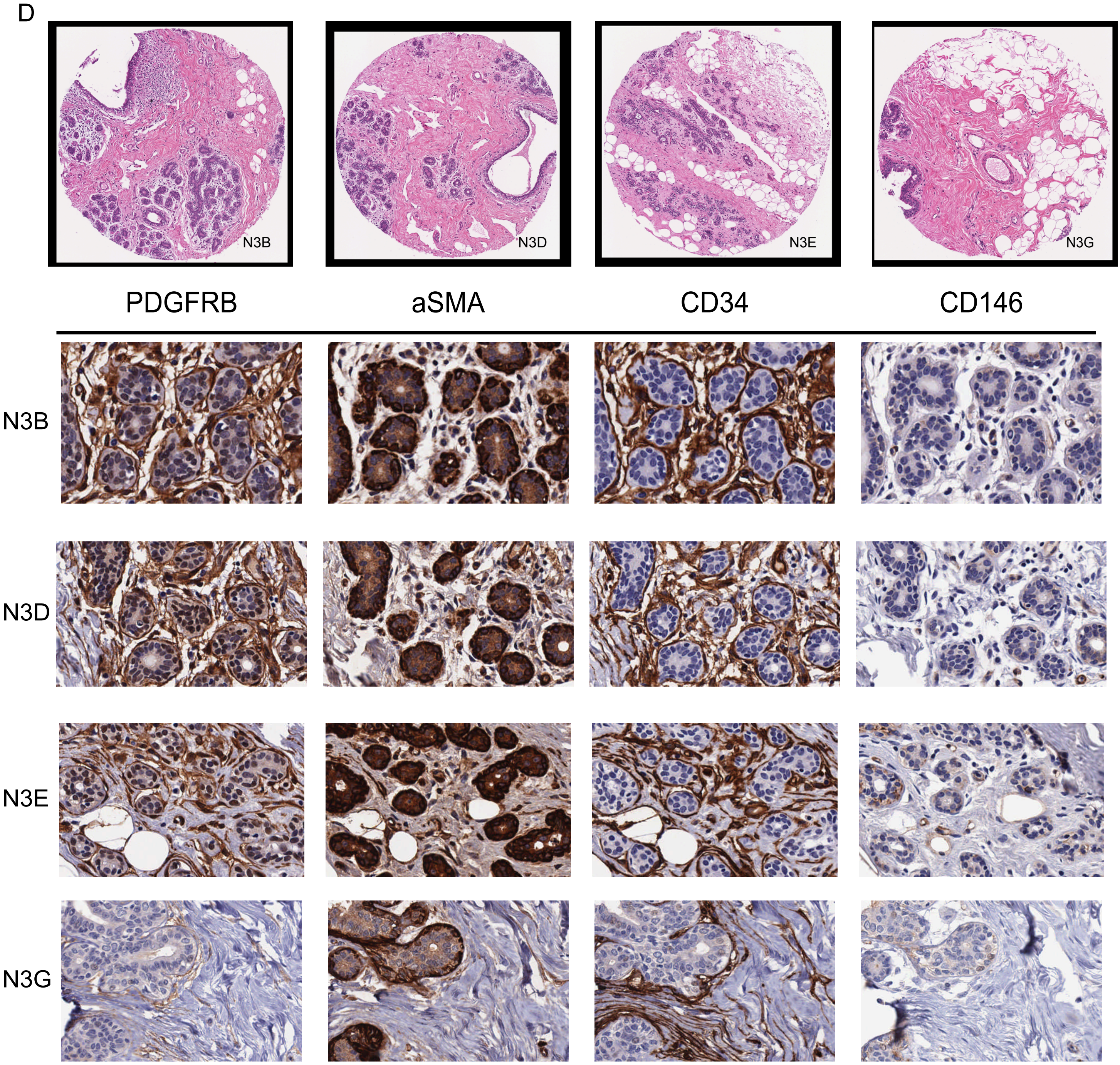


E

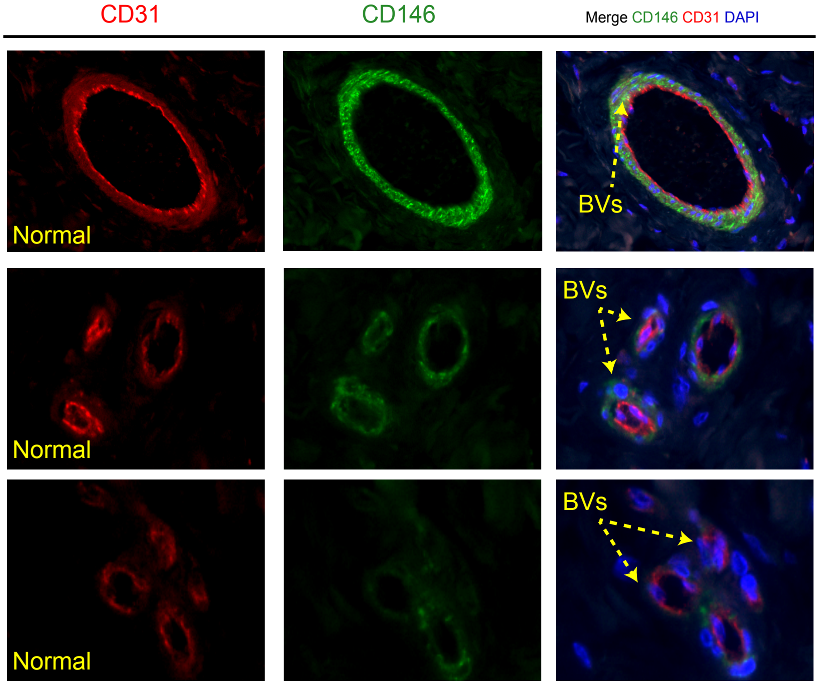

### Expanded View 5

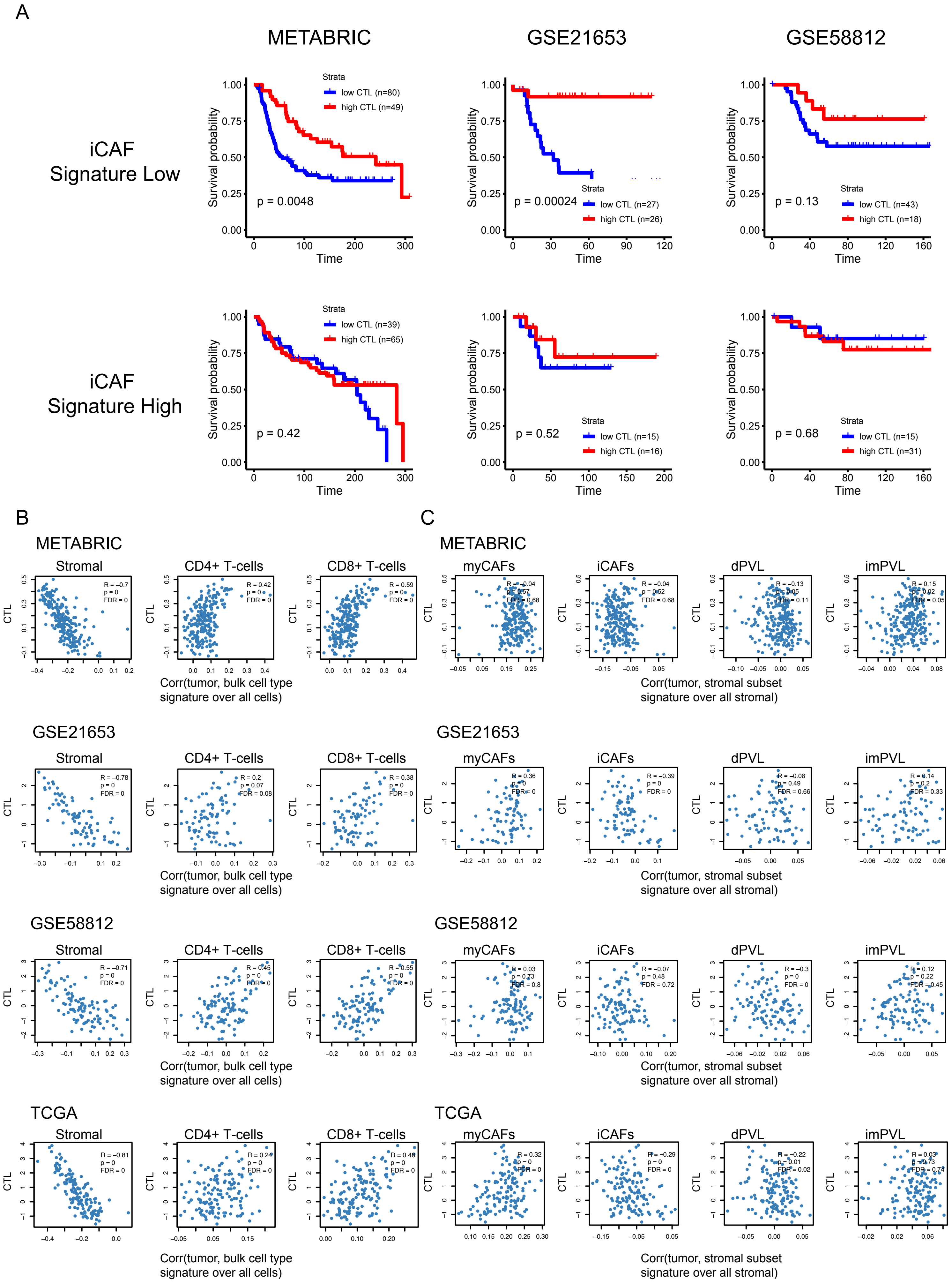
