## Appendix Figures for "Single-cell analysis reveals diverse stromal subsets associated with immune evasion in triple-negative breast cancer"

Appendix Figure S1

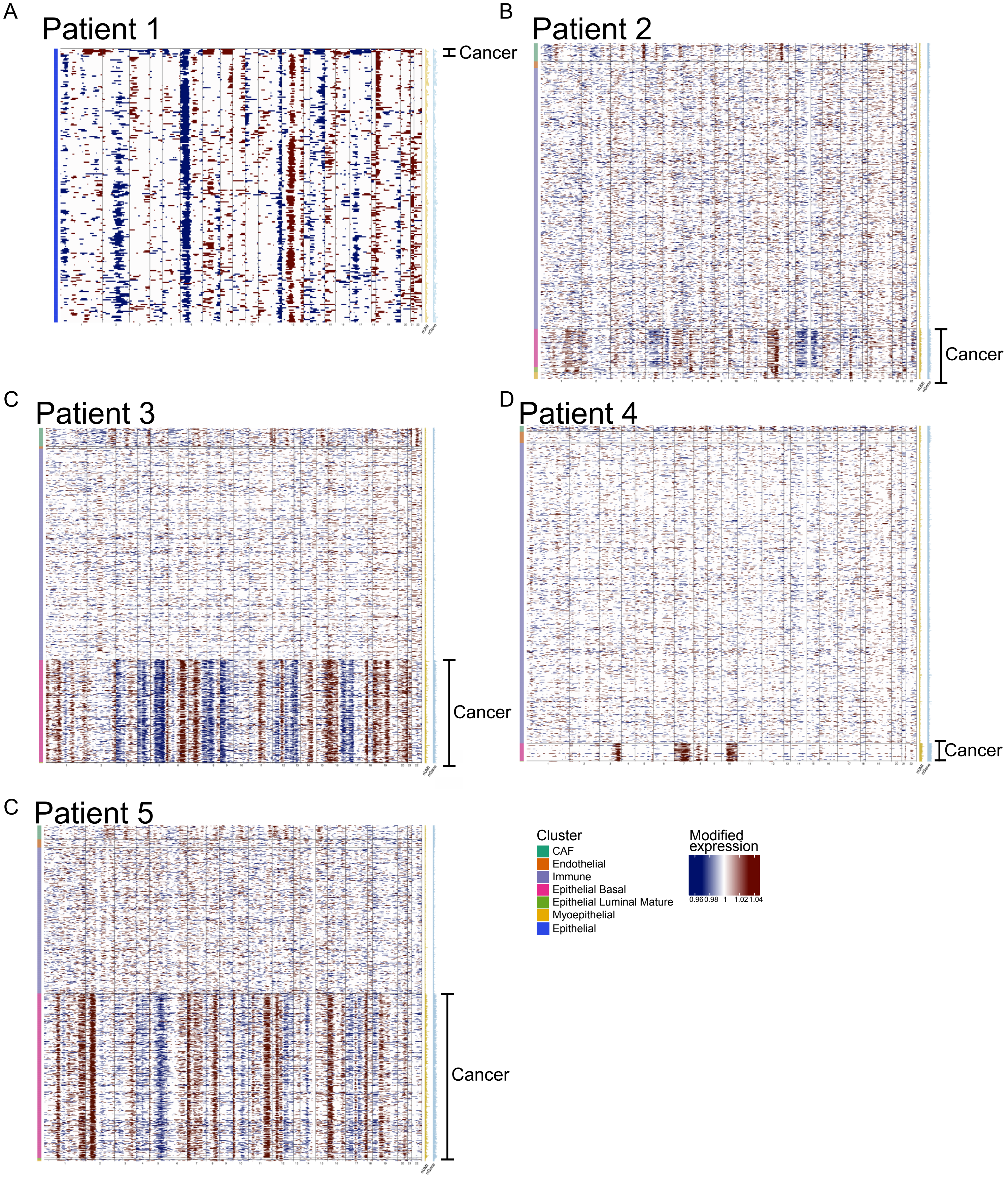

### Appendix Figure S2

A

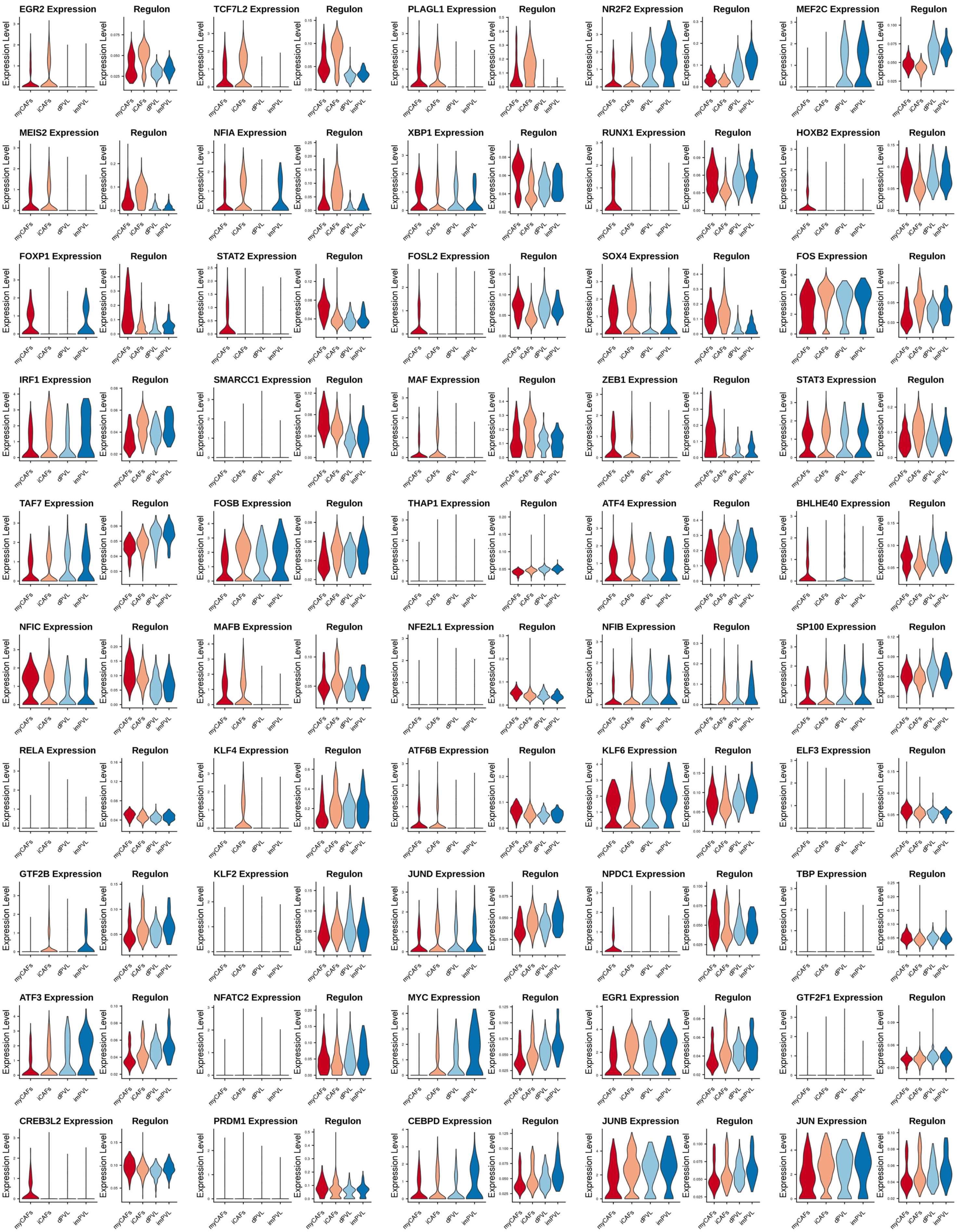

Appendix Figure S2 (cont.)

B

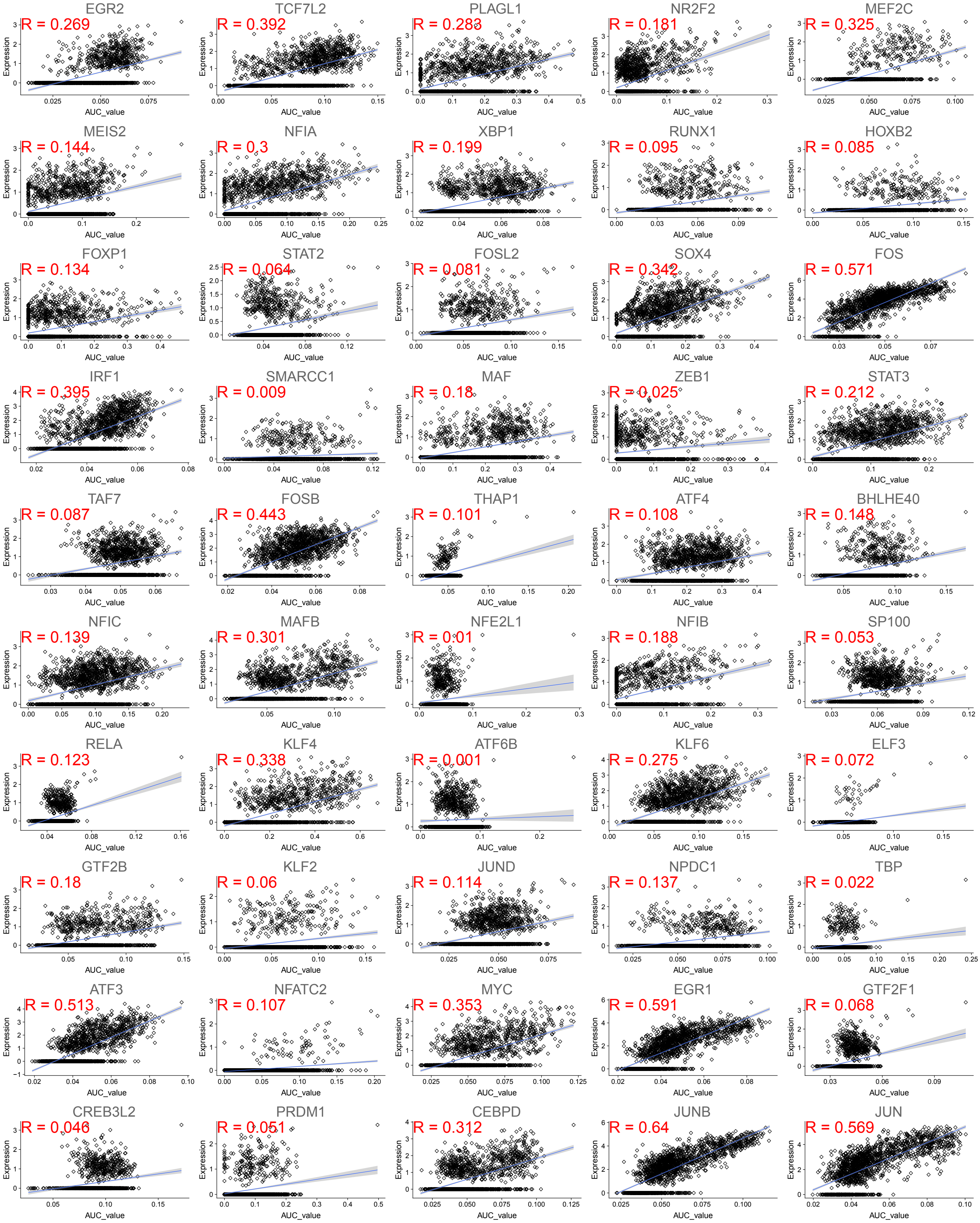

### Appendix Figure S3

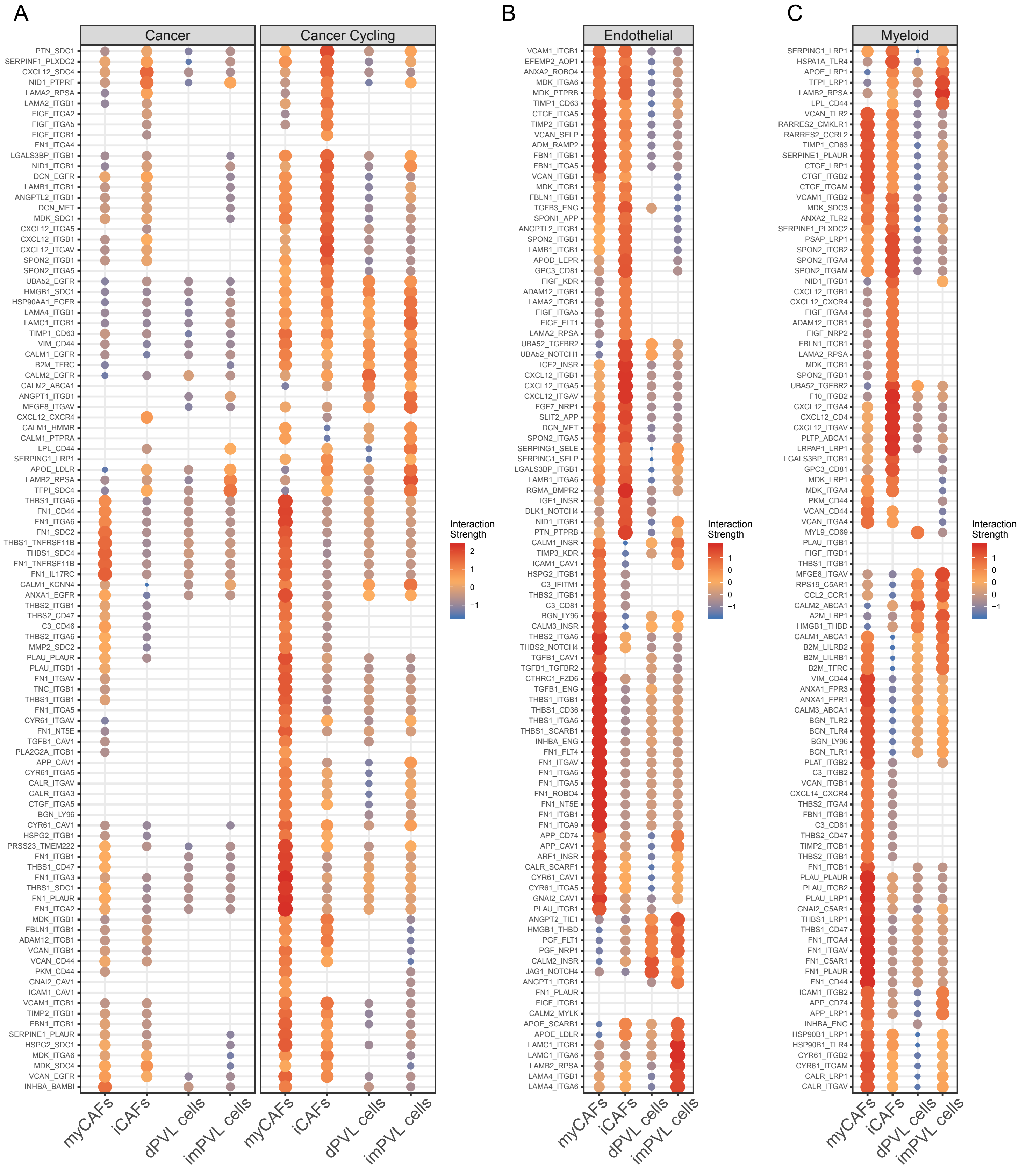

Appendix Figure S3 (cont.)

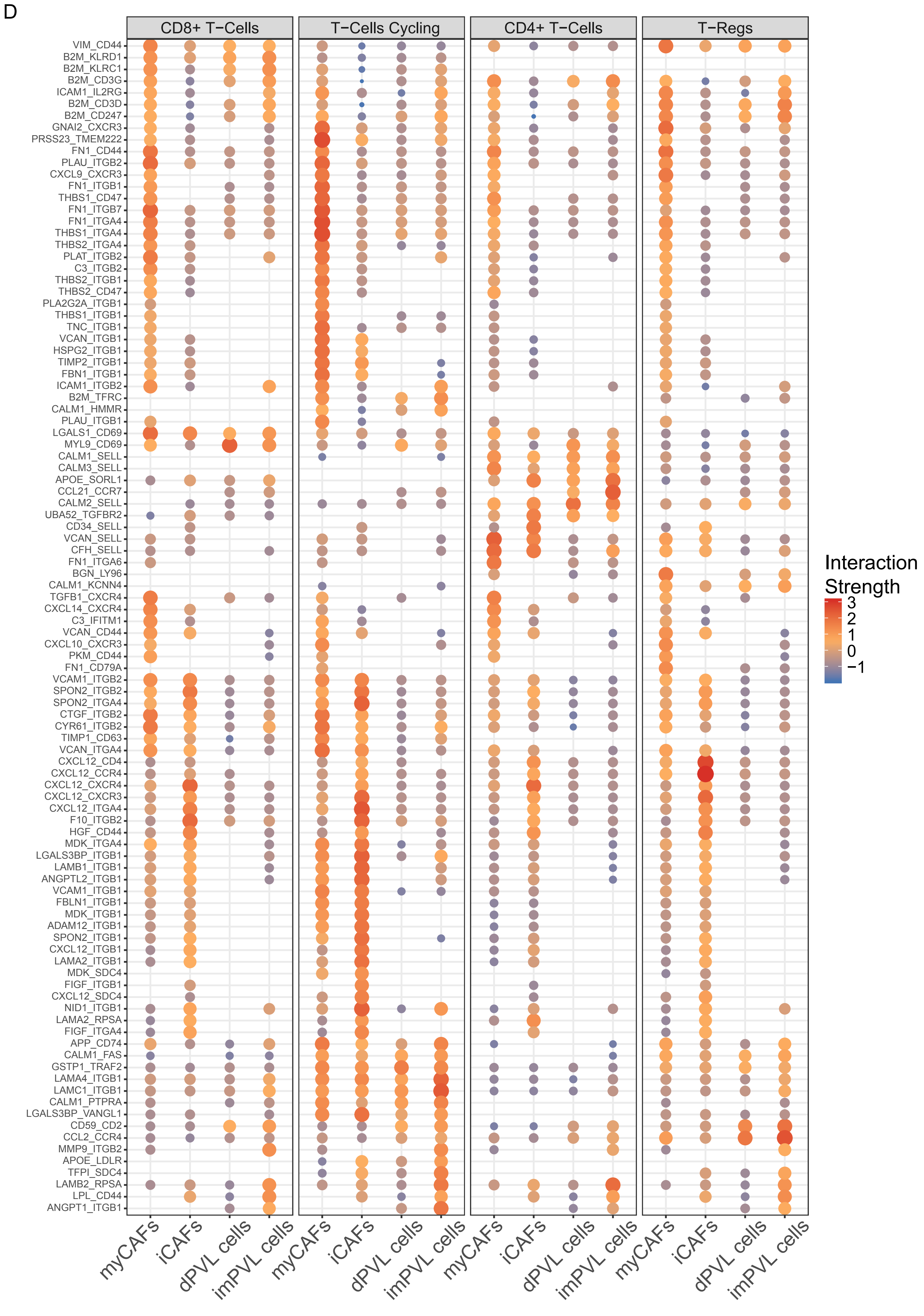

Appendix Figure S4

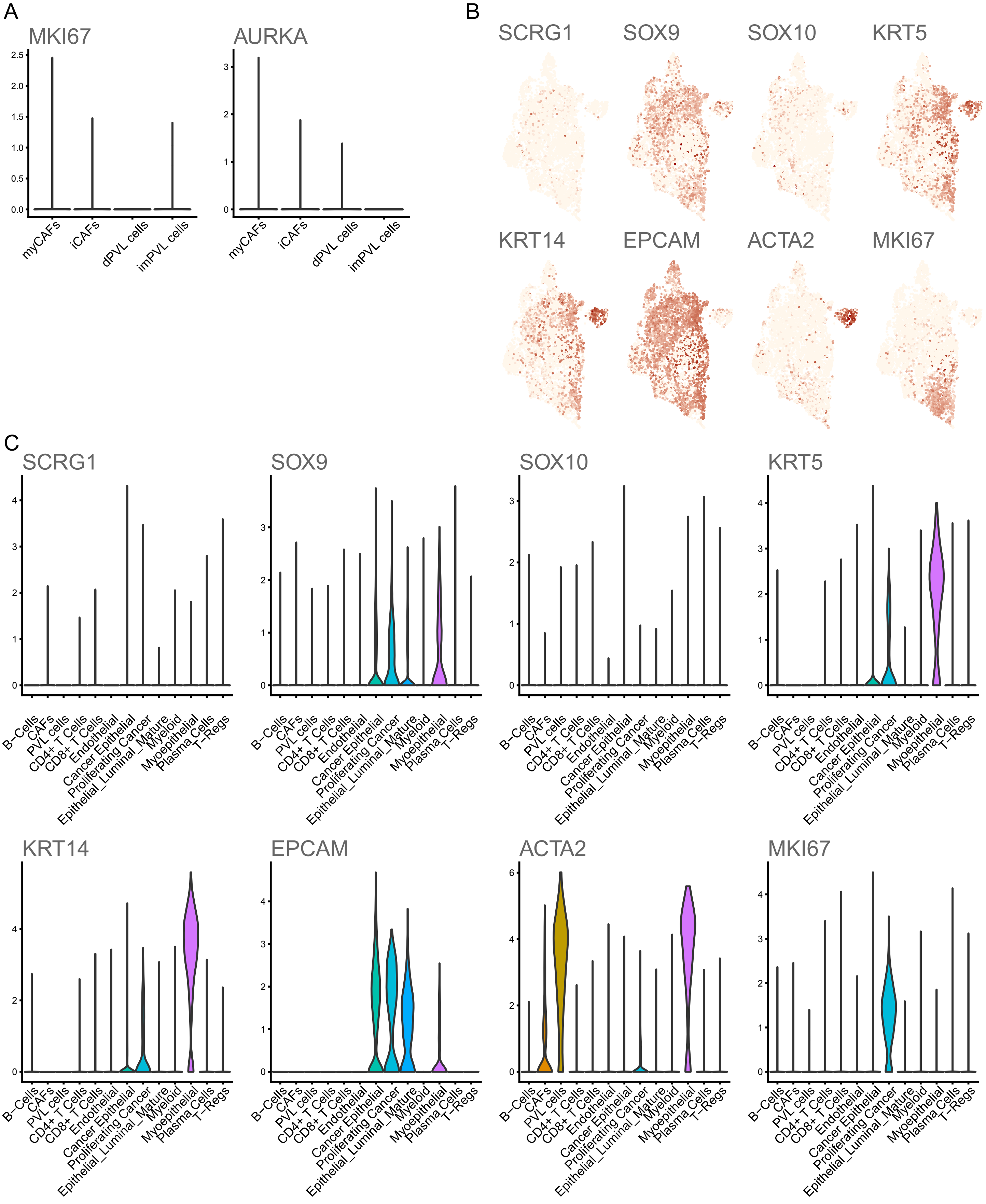
